# Systems medicine dissection of chromosome 1q amplification reveals oncogenic regulatory circuits and informs targeted therapy in cancer

**DOI:** 10.1101/2021.11.17.469031

**Authors:** Nikolaos Trasanidis, Alexia Katsarou, Kanagaraju Ponnusamy, Yao-An Shen, Ioannis V Kostopoulos, Bien Bergonia, Keren Keren, Paudel Reema, Xiaolin Xiao, Richard M Szydlo, Pierangela MR Sabbattini, Irene AG Roberts, Holger W Auner, Kikkeri N Naresh, Aristeidis Chaidos, Tian-Li Wang, Luca Magnani, Valentina S Caputo, Anastasios Karadimitris

## Abstract

Understanding the biological and clinical impact of copy number aberrations (CNA) in cancer remains an unmet challenge. Genetic amplification of chromosome 1q (chr1q-amp) is a major CNA conferring adverse prognosis in several cancers, including the blood cancer, multiple myeloma (MM). Although several chr1q genes portend high-risk MM disease, the underpinning molecular aetiology remains elusive. Here we integrate patient multi-omics datasets with genetic variables to identify 103 adverse prognosis genes in chr1q-amp MM. Amongst these, the transcription factor PBX1 is ectopically expressed by genetic amplification and epigenetic activation of its own preserved 3D regulatory domain. By binding to reprogrammed super-enhancers, PBX1 directly regulates critical oncogenic pathways, whilst in co-operation with FOXM1, activates a proliferative gene signature which predicts adverse prognosis across multiple cancers. Notably, pharmacological disruption of the PBX1-FOXM1 axis, including with a novel PBX1 inhibitor is selectively toxic against chr1q-amp cancer cells. Overall, our systems medicine approach successfully identifies CNA-driven oncogenic circuitries, links them to clinical phenotypes and proposes novel CNA-targeted therapy strategies in cancer.

**Significance:** We provide a comprehensive systems medicine strategy to unveil oncogenic circuitries and inform novel precision therapy decisions against CNA in cancer. This first clinical multi-omic analysis of chr1q-amp in MM identifies a central PBX1-FOXM1 regulatory axis driving high-risk prognosis, as a novel therapeutic target against chr1q-amp in cancer.

## Introduction

Genetic amplification of chr1q (chr1q-amp), one of the most frequent copy number aberrations (CNA), confers adverse prognosis in cancer (1–3). In multiple myeloma (MM), an incurable cancer of the B lineage plasma cells (PC), chr1q-amp is a secondary genetic event present in 30-40% of patients at diagnosis and is associated with adverse prognosis, high-burden proliferative disease and drug resistance (4–7).

Previous studies, often guided by low resolution methodologies (e.g., FISH against 1q21 locus(8)), identified several chr1q21 genes associated with adverse prognosis in MM, including the *CKS1B, PDKZ1, ILF2, ARNT, ADAR1 and IL6R* genes(9–13). However, genetic amplification that extends beyond chr1q21 has been reported in a small cohort of MM patients(14), raising the prospect that additional chr1q regions contribute to the biological profile and clinical impact of chr1q-amp. Further, how genetic amplification affects the 3D chromatin architecture of chr1q and influences biological processes that promote high risk disease is not known. Understanding these processes could inform novel anti-cancer therapeutic approaches targeted to chr1q-amp that are currently lacking.

Here we employed a comprehensive systems medicine approach to resolve the 3D genome landscape of chr1q-amp and to integrate it with multi-omic patient datasets. This approach led to the identification of adverse prognosis genes across the whole chr1q arm, and particularly in the 1q22 and 1q23.3 bands. Amongst 1q23.3-associated genes, we identified the transcription factor PBX1, which, in co-operation with FOXM1, regulates myeloma PC proliferation and generates a selective therapeutic vulnerability in chr1q-amp MM that can be targeted by a novel PBX1 inhibitor.

## Results

### Distinct patterns of amplification within chr1q shape its 3D chromatin architecture

We first explored whether and how genomic structural changes might impact the 3D chromatin structure of chr1q-amp. For this purpose, we constructed a correlation matrix of copy number scores across the chr1q arm (2D genome co-amplification map) using WGS data from MM patients (MMRF database (15), n=896) and compared it with 3D genome Hi-C contact maps of two chr1q-amp MM cell lines (MMCL; U266, RPMI8226 (16) ; **Fig. 1A**). By applying the same computational method used for topologically associated domain (TAD) discovery (16), we found four main blocks of co-amplification (termed topologically co-amplified domains; TCDs), which define distinct amplification patterns across MM patients (**Fig. 1A and Supplementary Fig. S1A**). Comparison of insulation score profiles across chr1q revealed poor correlation between 2D (WGS) and 3D (Hi-C) genome maps (**Supplementary Fig. S1B**). Additional analysis using Hi-C data from non-amplified, reference B-lineage cells (GM12828)(17) showed almost 65% of its TADs to be disrupted by chr1q-amp breakpoints (**Supplementary Fig. S1C**), suggesting that genetic amplification extensively disrupts the 3D chromatin architecture of chr1q. Nevertheless, we detected four large segments (B1-B4 hyper-domains) with overlapping TCD/TAD borders, suggesting the presence of amplification patterns that preferentially retain the overall chromatin structure of these four hyper-domains (**Supplementary Fig. S1D).**

**Fig. 1.**
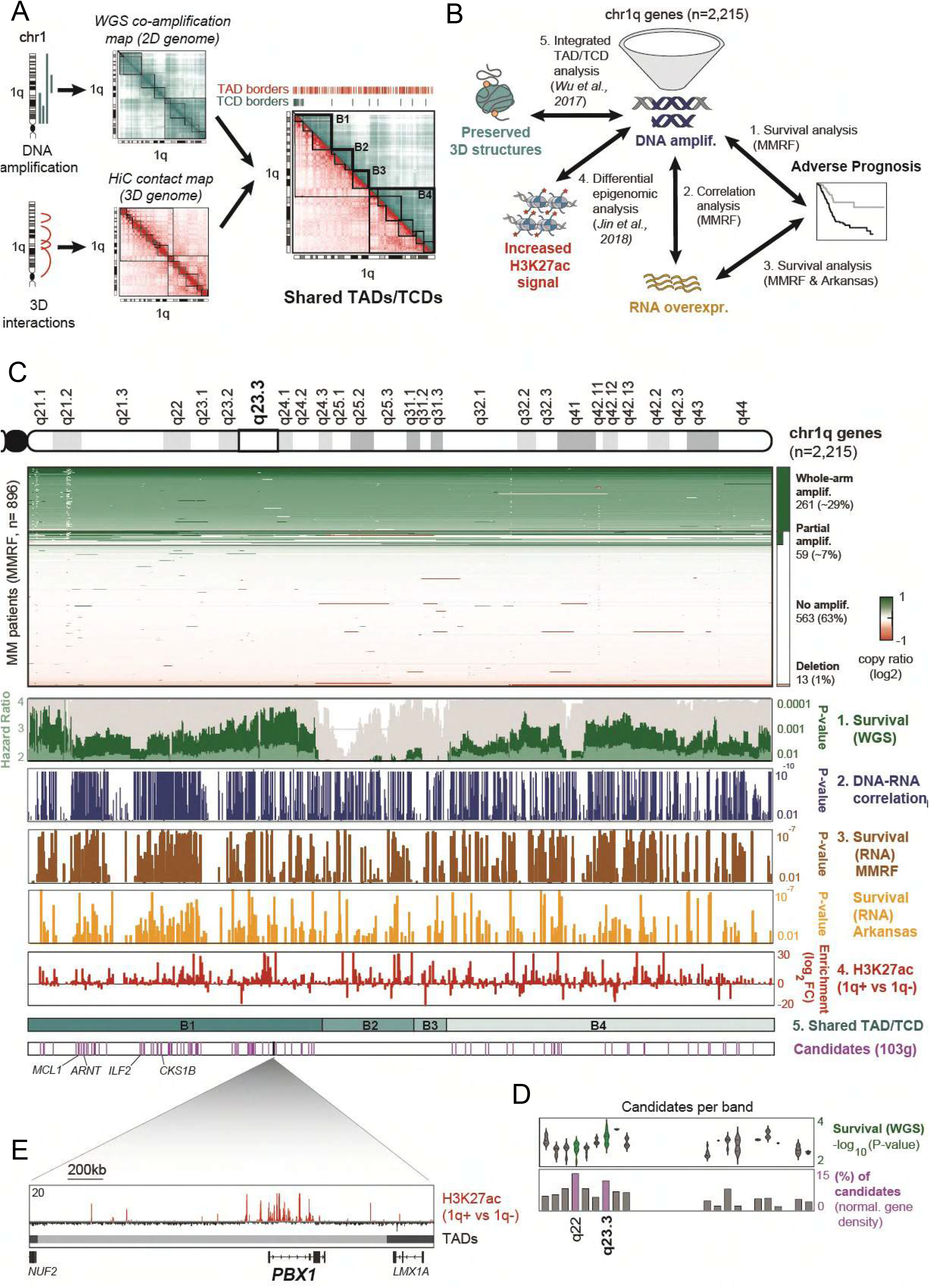
Multi-layer, systems medicine analysis of chr1q amplification in multiple myeloma. **(A)** Two-dimensional (cyan) co-amplification and three-dimensional (red) Hi-C contact maps of chr1q locus in MM cells used to identify topologically co-amplified domains (TCDs) and topologically associated domains (TADs), respectively. Map overlay identified four major co-amplified domains that retain a preserved 3D structure (B1-4 hyper-domains). **(B)** Schematic overview of the analysis strategy to detect candidate gene drivers of biological and clinical impact. Scanning across the chr1q locus (2,215 genes), genes fulfilling all criteria were considered as candidate drivers: (1) genetic amplification is significantly associated with poor prognosis (MMRF dataset, n=896); (2) genetic amplification is significantly associated with overexpression (MMRF dataset, n=896); (3) overexpression is associated with poor prognosis in MMRF dataset (top panel, n=896) and Arkansas dataset (bottom panel, n=413); (4) significant epigenetic activation (H3K27ac gain) is detected in chr1q-amplified versus non-amplified samples (Jin et al., n=10). The B1-B4 hyper-domains were also used as a reference here (5). **(C)** Analysis overview, from top to bottom: chr1q cytogenetic map; copy-number profiles of chr1q genes across MMRF patients detecting whole-arm amplification (~29%), partial amplification (~7%), no amplification (~63%) and deletions (~1%); survival analysis of genetic amplification of chr1q genes across MMRF patients (WGS, 73 genetic parameters; dark green bars, P-value; light green bars, Hazard Ratio; grey bars, % bootstrapping confidence levels); Pearson correlation analysis between copynumber ratios (WGS) and expression (RNA-seq; blue bars indicate Pearson correlation p-values); survival analysis of chr1q gene expression (RNA-seq) in MMRF (brown) and Arksansas (yellow) datasets (bars indicate analysis p-values); differential H3K27ac analysis between chr1q-amplified (n=5) versus non-amplified (n=5) MM cells (red bars indicate differential log2 fold-change enrichment scores); four chr1q domains (B1-4) with conserved TAD/TCD structures; Candidate pathogenic driver genes (n=103, pink bars) identified by the current analysis (the previously known *MCL1, ARNT, ILF2* and *CKS1B* genes are shown here). **(D)** Analysis overview of candidate driver genes (103) across chr1q bands. Distribution of WGS multivariate analysis scores (-log10P-value; top) and percentage (%) of candidate genes (relative to band gene density) per cytogenetic band. The highest candidate genes density was detected in 1q22 and 1q23.3 bands (highlighted here), with 1q23.3 also displaying the highest survival significance scores. **(E)** The *PBX1* gene as a prominent candidate occupying alone a single TAD, displays strong epigenetic activation across PBX1 body and putative enhancers in chr1amp myeloma PC.

### Systems medicine analysis identifies adverse prognosis drivers beyond 1q21

Next, to identify genes across chr1q that could drive high-risk phenotype in MM and with reference to the resolved 3D chromatin structure, we combined genomic (WGS, WES), epigenomic (H3K27ac-seq), and transcriptomic (RNA-seq, DNA microarrays) data with genetic variables from three previous studies: MMRF (n= 896); Arkansas (n=414); and *Jin2018* (n=12) (15, 16, 18, 19) (**Fig. 1B**). Of the 2,215 chr1q genes, we considered as candidate drivers only genes fulfilling each of the following criteria: (1) their genetic amplification predicts adverse prognosis, independent of the prognostic impact of 73 other molecular markers (MMRF dataset; **Supplementary Fig. S1E**); (2) their genetic amplification is significantly associated with their transcriptional overexpression (MMRF dataset); (3) their overexpression is significantly correlated with adverse prognosis (MMRF and Arkansas datasets); (4) their genetic amplification is accompanied by epigenetic activation (i.e., H3K27ac signal gain compared to non-amplified MM; *Jin2018* dataset); (**Fig. 1C and Supplementary Table S1**).

This stepwise analysis identified 103 candidate genes residing exclusively in B1 and B4 hyper-domains, including the previously known *MCL1, CKS1B, ILF2* and *ARNT* genes in chr1q21.3 (9–11) (**Fig. 1C**). Pathway analysis of all 103 genes showed significant enrichment for cell cycle-related processes, suggesting their direct involvement in the proliferative phenotype that is associated with chr1q-amp MM (13) (**Supplementary Fig. S1F**). Interestingly, we identified two cytogenetic bands, 1q22 and 1q23.3, to contain the highest number of candidate adverse prognosis genes, relative to their gene density (**Supplementary Fig. S1G**), with 1q23.3 displaying the highest association to adverse prognosis (**Fig. 1C and 1D**). Therefore, there are additional regions, other than 1q21, which contribute to the high-risk, proliferative phenotype linked to chr1-amp in MM.

### PBX1 is a novel biomarker of chr1q genetic amplification

Amongst 1q23.3 genes, the transcription factor *PBX1* previously reported to promote cancer cell survival, metastasis and drug resistance (20–22) was notable for the highest H3K27ac signal gain across its own preserved TAD (**Fig. 1E and Supplementary Fig. S1D and S1H**). These features comprise a unique case of amplification of an entire regulatory domain linked to epigenetic activation, gene overexpression and adverse prognosis. Further analysis using the MMRF dataset confirmed *PBX1* as a marker of high-risk MM disease, with its amplification significantly correlating with its overexpression (**Supplementary Fig. S2A and S2B**), while *PBX1* overexpression was associated with high-risk clinical features, high myeloma plasma cell proliferative index, progressive/relapsed disease and worse overall survival (**Supplementary Fig. S2C-S2J**).

### The pro-proliferative role of PBX1 in chr1q-amp MM

We explored further the functional role of *PBX1* in chr1q-amplified MM cells, by assessing its mRNA and protein expression levels across healthy and tumour cells. We found that in normal hematopoiesis, *PBX1* is expressed in bone marrow hematopoietic stem and progenitor cells as well as megakaryocytes, but not in B cells or plasma cells (**Supplementary Fig. S3A**). In MM, we confirmed ectopic expression of *PBX1* in four chr1q-amp MMCL (**Fig. 2A**) and in 9/11 patient myeloma PC samples with FISH-verified chr1q-amp (**Fig. 2B and Supplementary Fig. S3B and S3C**).

**Fig. 2.**
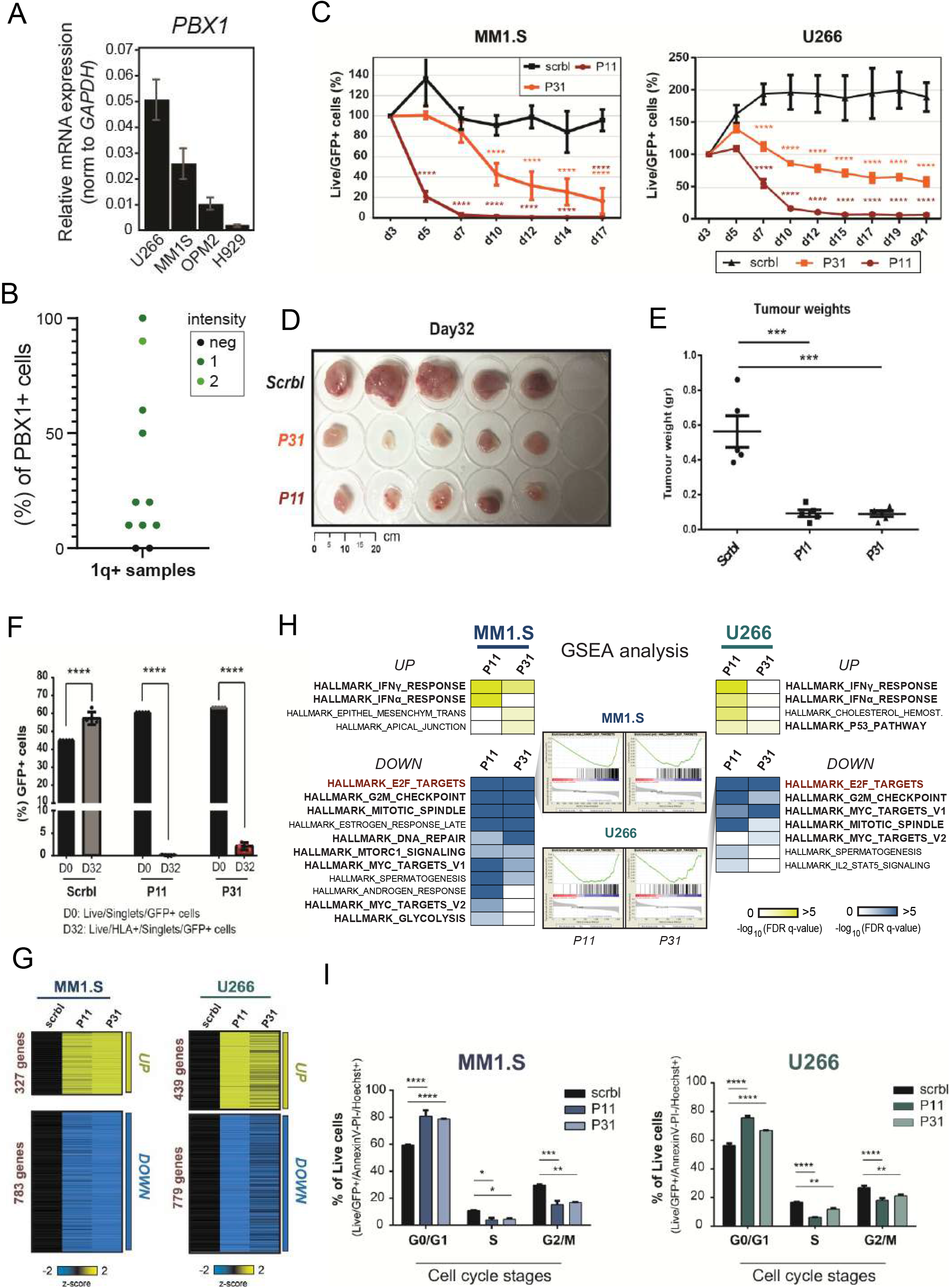
PBX1-dependent myeloma cell proliferation. **(A)** mRNA expression of PBX1 in four MM cell lines. **(B)** Immunohistochemical (IHC) analysis of trephine bone marrow samples from 11 MM patients detects no (neg), medium (1) or high (2) PBX1 expression at clonal or subclonal level (% of PBX1+ cells). **(C)** Time-course, flow-cytometry based analysis of MM1.S (left) and U226 (right) myeloma cell viability *in vitro,* upon lentiviral transduction with scrambled control (scrbl) and anti-PBX1 shRNAs (P11, P31). Data collected from three biological replicates represent the fraction of GFP+ live cells on the timepoints displayed, after normalization against Day3. Statistical analysis was performed using a twoway ANOVA with post-hoc multiple comparisons test. Error bars represent SEM (n=3). **(D & E)** Knock-down of PBX1 in MM1.S cells using an *in vivo* plasmacytoma xenograft mouse model; tumour size photograph **(D)** and tumour weights **(E)** measured at termination date (Day 32). Statistical analysis performed using Kruskall-Wallis with Dunn’s post-hoc multiple comparisons test. **(F)** Relative fraction of transduced cells detected at start (Day0, Live/GFP+ cells) and termination (Day32, Live/HLA+GFP+ cells) dates. **(G)** RNA-seq analysis of PBX1-depleted MM1.S and U266 cells 3 days after lentiviral transduction. Heatmaps indicate differentially expressed genes shared between P11- and P31-depleted cells for each cell line. **(H)** Gene Set Enrichment Analysis (GSEA) of up- (top) or down-regulated (bottom) genes in MM1.S (left) and U266 (right) myeloma cells illustrating significantly enriched molecular pathways in each cell line. Enrichment plots for the prominent cell cycle regulation pathway (E2F targets), which was identified as a top hit, are also presented here. **(I)** Flow-cytometric cell-cycle analysis of MM1.S and U266 cells 6 days after PBX1 knockdown. Data present the summary of 3 biological experiments. Analysis was done using parametric one-way ANOVA with post-hoc multiple comparisons test. *\*:P<0.05;**:P<0.01; ***:P<0.001; ****:P<0.0001*

Depletion of *PBX1* using two validated shRNAs (P31, P11) and assessed by GFP marker expression was toxic to MM1.S and U266 cells compared to scrambled shRNA control *in vitro* (**Fig. 2C and Supplementary Fig. S3D**) and impaired myeloma cell growth (MM1.S) in an *in vivo* subcutaneous MM model **(Fig. 2D-2F and Supplementary Fig. S3E-S3G)**. To gain further insights, we performed RNA-seq analysis in both MMCL upon shRNA-mediated PBX1 depletion (**Fig. 2G-2H and Supplementary Table S2**). Transcriptome profiling of *PBX1*-depleted cells showed similar numbers of genes de-regulated in the two MMCL, while Gene Set Enrichment Analysis revealed significant enrichment for cell cycle-related pathways amongst down-regulated and interferon response pathways in up-regulated genes **(Fig. 2H).** This is consistent with the reported enrichment of interferon response pathways in early-stage, non-proliferative MM and of cell cycle-related pathways in advanced disease and MMCL (23, 24). Accordingly, flow-cytometric analysis showed significant G1-phase cell cycle arrest in *PBX1-* depleted MMCL **(Fig. 2I, Supplementary Fig. S3H).**

### Defining the epigenetic and regulatory programme of PBX1 in chr1q-amp cells

ChIP-seq analysis against PBX1 in MM1.S and U266 cells identified 30,000-40,000 binding sites (**Fig. 3A and Supplementary Table S2**). Further annotation using chromHMM maps (built upon ENCODE/Blueprint Consortium data) showed that 60-80% of PBX1 recruitment occurs in active-chromatin promoter and enhancer areas, while motif enrichment analysis identified the PBX1 motif among the top hits **(Fig. 3A and Supplementary Fig. S4A-S4D)**. Additional analysis of H3K27ac-seq profiles from eight primary myeloma PC and nine MMCL(19) identified 2,400 super-enhancers (SEs), 70% of which are PBX1-bound **(Fig. 3B)**. Samples stratification based on chr1q-amp status showed significantly higher H3K27ac signal in PBX1-bound SEs in chr1q-amplified versus non-amplified cells, suggesting extensive epigenetic reprogramming associated with PBX1 binding in chr1q-amplified myeloma cells **(Fig. 3C and Supplementary Fig. S4E-S4F)**. Interestingly, the PBX1-bound SEs in chr1q-amplified cells are predicted to regulate critical cellular pathways, including cell cycle (**Fig. 3D**).

**Fig. 3.**
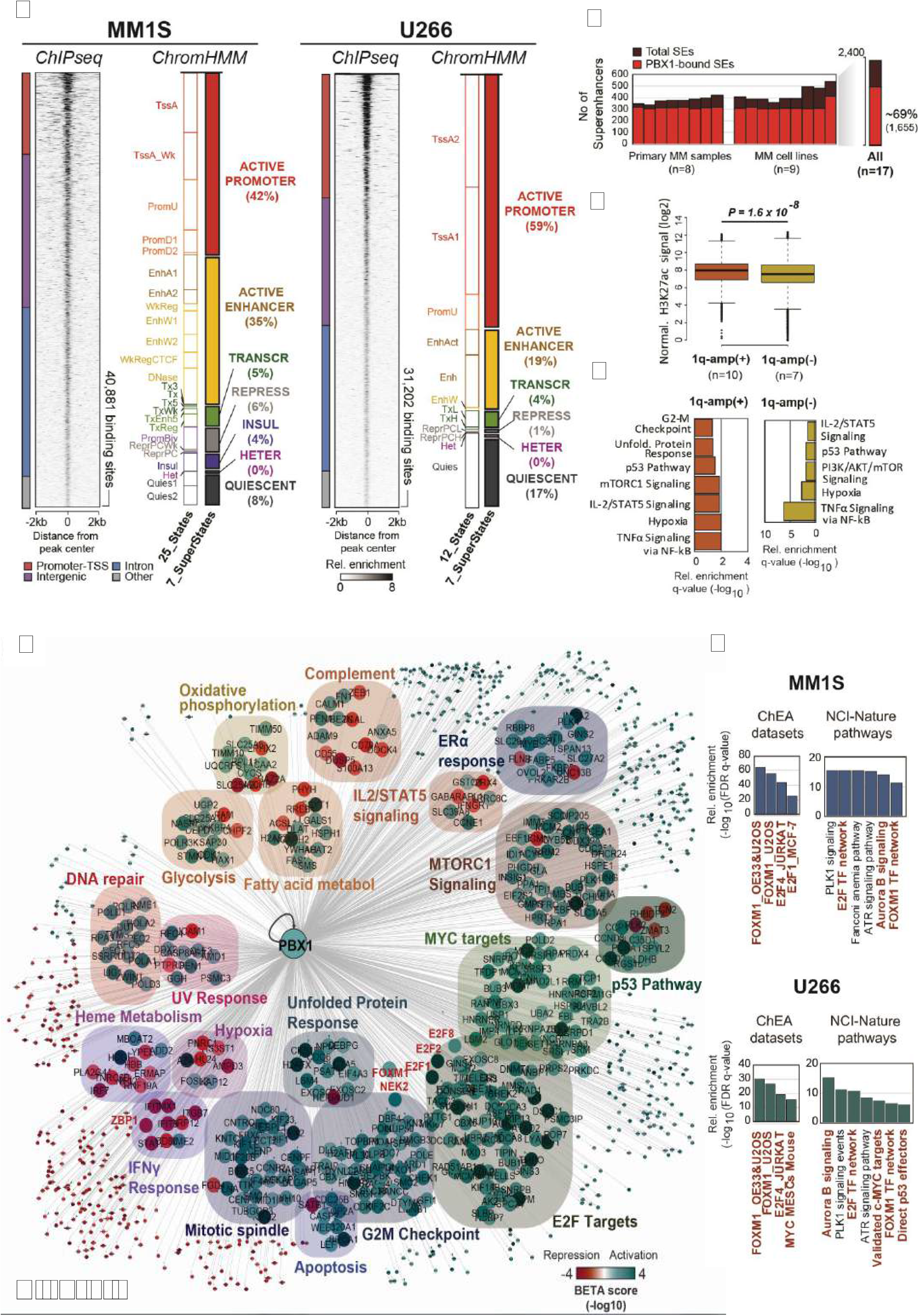
Genome-wide analysis of PBX1 function in chr1q-amplified myeloma cells. **(A)** Heatmap representation of PBX1 cistrome in MM1.S and U266 cells, as identified by ChIP-seq analysis (n=2 per cell line). Genomic annotation (left) and epigenomic chromHMM states (right) of significantly enriched regions are also presented here. **(B)** Super-enhancer (SEs) analysis across 9 MM cell lines and 8 MM primary samples using H3K27ac ChIP-seq (data obtained from *Jin et al., 2018).* Number of total (dark red) and PBX1-bound (red) SEs (red) across 17 MM samples and aggregated profile in all samples (right) is shown. **(C)** Boxplot representations of average normalized H3K27ac signal of chr1q-amplified and nonamplified samples across 1,655 PBX1-bound SEs. Analysis was performed using Mann-Whitney t-test. **(D)** Pathway analysis of genes predicted to be regulated by PBX1-bound SEs in chr1q-amplified (+) and non-amplified (-) cells. **(E)** Integrative cistrome-transcriptome analysis with BETA-plus displays the regulatory programme of PBX1 in MM1.S cells. Biological annotation of genes was performed using the Molecular Signatures Database. Node colours represent average predicted activation (blue) or repression (red) for each gene. Transcriptional targets of interest are highlighted in red font. **(F)** Overrepresentation analysis against the ChEA database and NCI-Nature pathways of the direct PBX1 target genes in MM1.S (top) and U266 (bottom) cells. Terms of interest are highlighted in red font.

Next, we integrated the PBX1 cistrome with the PBX1-depleted transcriptomes to generate the gene regulatory network of PBX1 in chr1q-amplified cells (**Fig. 3E and Supplementary Fig. S4G-S4I and Supplementary Table S3**). We identified approximately 700 and 300 genes to be directly activated and repressed, respectively, by PBX1 in both MM1.S and U266 MMCL. Again, among other prominent oncogenic pathways, the former were primarily enriched in cell cycle-related biological processes and the latter in interferon response pathways (**Fig. 3E**).

### The PBX1-FOXM1 axis regulates cell proliferation in chr1q-amp MM

Amongst the PBX1-dependent targets, we detected significant enrichment of the pro-proliferative *FOXM1* and *E2F* transcription factors and their corresponding targets (**Fig. 3F**), such as the FOXM1-dependent *NEK2* that regulates drug resistance in MM (25, 26), **(Fig. 4A)**. Further, we identified PBX1 binding on active *PBX1, E2F1/2, NEK2* promoters and *PBX1, FOXM1, E2F2, NEK2* enhancers (**Fig. 4B**), while FOXM1 was found to bind to the same *FOXM1* and *NEK2* regions as PBX1 (**Supplementary Fig. Fig S5A**). To better explore the regulatory interplay among those factors (**Fig. 4A**), we characterized further the role of FOXM1 in chr1q-amp cells. Knockdown of *FOXM1* using two validated shRNAs was toxic to MM1.S cells **(Fig. 4C)**, as previously shown ^(25)^. In addition, depletion of *FOXM1* mRNA was associated with downregulation of *NEK2* but not of *PBX1* (**Fig. 4D**), suggesting that *FOXM1* acts downstream of *PBX1* (**Fig. 4A**). Moreover, RNAseq analysis revealed approximately 800 differentially expressed genes after *FOXM1* knockdown in MM1.S cells (**Fig. 4E**), with cell cycle-related pathways found to be significantly enriched amongst downregulated genes **(Fig. 4F).** Cell cycle arrest at G2/M was corroborated by flow-cytometry, thus confirming the pro-proliferative role of FOXM1 in chr1q-amplified MMCL **(Supplementary Fig. S5B)**.

**Fig. 4.**
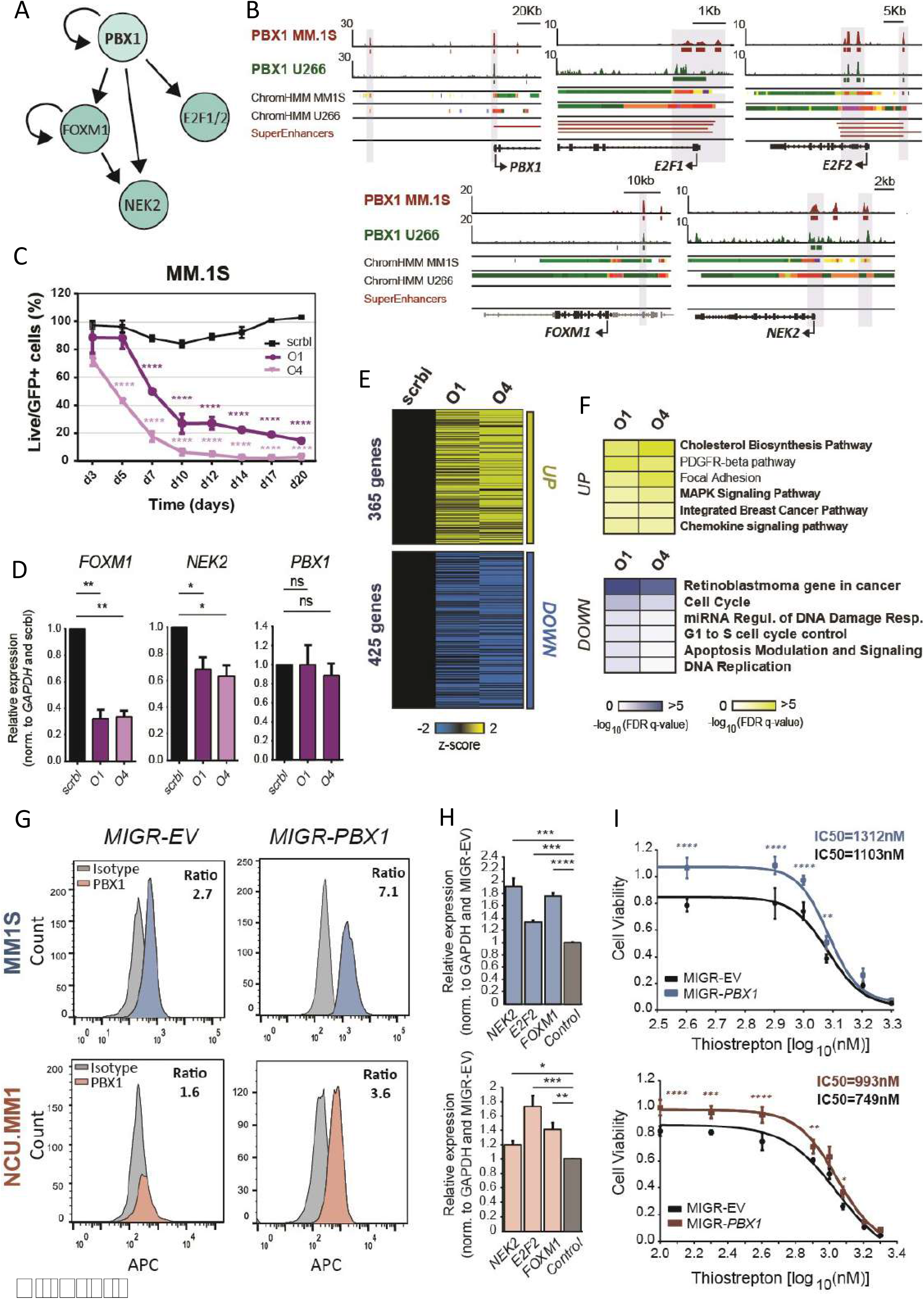
PBX1 regulates directly FOXM1- and E2F1/2-associated transcriptional programmes in chr1q-amplified MM cells. **(A)** Regulatory connections between PBX1 and its downstream targets FOXM1, E2F1/2 and NEK2 in chr1q-amplified MM cells as emerged from data shown in b-i. **(B)** IGV snapshots display the epigenomic features of prominent genetic loci: *PBX1* promoter and enhancer*, E2F1* promoter*, E2F2* promoter and enhancer*, FOXM1* enhancer*, NEK2* promoter and enhancer. From top to bottom: PBX1 ChIP-seq in MM1.S and U266 cells; ChromHMM maps in MM1.S and U266 cells (colour code same as **Fig 3A)**; Super-enhancers are as identified in chr1q-amplified MMCL and primary samples. **(C)** Flow cytometry-based analysis of MM1.S cells survival (n=3) upon transduction with anti-*FOXM1* shRNAs (O1, O4) and scrambled control (scrbl) lentiviral vectors. Statistical analysis was performed by a two-way ANOVA with post-hoc multiple comparisons test. **(D)** Analysis of PBX1, *FOXM1* and *NEK2* expression levels by RT-qPCR after lentiviral transduction with anti-*FOXM1* and scrambled control shRNA in MM1.S cells (n=3). Statistical analysis was performed using a one-way ANOVA with post-hoc multiple comparisons test. **(E)** Heatmap representation of differentially expressed genes after *FOXM1* depletion with O1 and O4 shRNAs in comparison to scrambled control (RNA-seq, n=2). **(F)** Over-representation analysis of significantly upregulated (top) and downregulated (bottom) genes upon *FOXM1* knockdown in MM1.S cells. **(G)** Intracellular staining followed by flow-cytometric analysis of MM1.S (top) and NCU.MM1 (bottom) cells transduced with control (MIGR-EV) or PBX1-overexpressing (MIGR-PBX1) vectors using anti-PBX1 or isotype control antibodies (mean fluorescence intensity ratio between antibodies is shown). **(H)** RT-qPCR analysis of *NEK2, E2F2 and FOXM1* mRNA expression in PBX1-overexpressing versus control MM1.S (top) and NCUMM1 (bottom) cells (n=4). Data were analysed using a one-way ANOVA with post-hoc multiple comparisons test. **(I)** Drug sensitivity assays in MIGR-EV and MIGR-*PBX1* transduced MM1.S (top) and NCU.MM1 (bottom) cells 48h after treatment with the FOXM1 inhibitor, thiostrepton (n=3). IC_50_ values were calculated for each cell line using a non-linear fitting model (fitting line represented here). Error bars show standard errors of mean *: *P<0.05;* **: *P<0.01;* ***: *P<0.001;* ****: *P<0.0001; n/s: not significant.*

For further validation of the PBX1-FOXM1 axis, we forced expression of exogenous *PBX1* into MM1.S and NCU.MM1 chr1q-amplified MM cells (**Fig. 4G**). This led to modest but significant increase in *FOXM1, NEK2* and *E2F2* mRNA levels (**Fig. 4H**) and significantly reduced sensitivity of the MMCL to thiostrepton, an inhibitor of FOXM1 transcription (25, 27) (**Fig. 4I and Supplementary Fig. S5C**). Rescue of *PBX1* depletion by shRNA-resistant *PBX1* cDNA resulted in a significantly lower MMCL toxicity, ameliorated cell cycle arrest and dampened downregulation of *FOXM1, NEK2* and *E2F2* (**Supplementary Fig. S5D-5G**), thus validating the genetic and functional interactions in the PBX1-FOXM1 (**Fig. 4A**) axis and its role in orchestrating an oncogenic, proliferative process in chr1q-amp MM cells.

### The PBX1-FOXM1 regulatory axis generates a selective therapeutic vulnerability in primary chr1q-amp MM cells

Next, we sought to validate activity of the PBX1-FOXM1 axis in primary myeloma plasma cells (**Fig. 5A**). For this purpose, we combined RNA-seq with ATAC-seq profiling of highly purified chr1q-amplified (n=6) and non-amplified (n=6) primary myeloma PC, and explored differences in chromatin accessibility, gene expression and predicted TF connectivity (**Fig. 5A and Supplementary Table S4**). In addition to previously established gene-markers (*CKS1B, IL6R, ARNT, PDKZ1, ADAR*), we also found overexpression of all main PBX1-FOXM1 module components (*PBX1, FOXM1, E2F1/2, NEK2*) in chr1-amp cells (**Fig. 5B**). Moreover, there was significant enrichment of proliferative pathways and FOXM1-dependent targets in genes overexpressed in chr1q-amp cells (**Fig.5C**). Comparative ATAC-seq analysis revealed enhanced chromatin accessibility in the regulatory regions of genes over-expressed in the same cells (**Fig. 5D**). Differential TF footprinting analysis revealed a higher number of TFs with increased connectivity (measured as differential regulatory potential, *ΔP*) in chr1q-amp versus nonamplified cells (**Fig. 5E)**. By combining transcriptional and regulation profiles, we identified 34 TFs with increased expression and connectivity in chr1q-amplified cells, including all four TFs involved in the PBX1-FOXM1 module (PBX1, FOXM1, E2F1, E2F2; **Fig. 4A and 5F).** Notably, as compared to nonamplified cells (n=3), chr1q-amplified primary myeloma cells (n=3) were selectively sensitive to thiostrepton treatment, while expression of *FOXM1* and *NEK2,* but not *PBX1,* decreased in response to treatment (**Fig. 5G and 5H**).

**Fig. 5.**
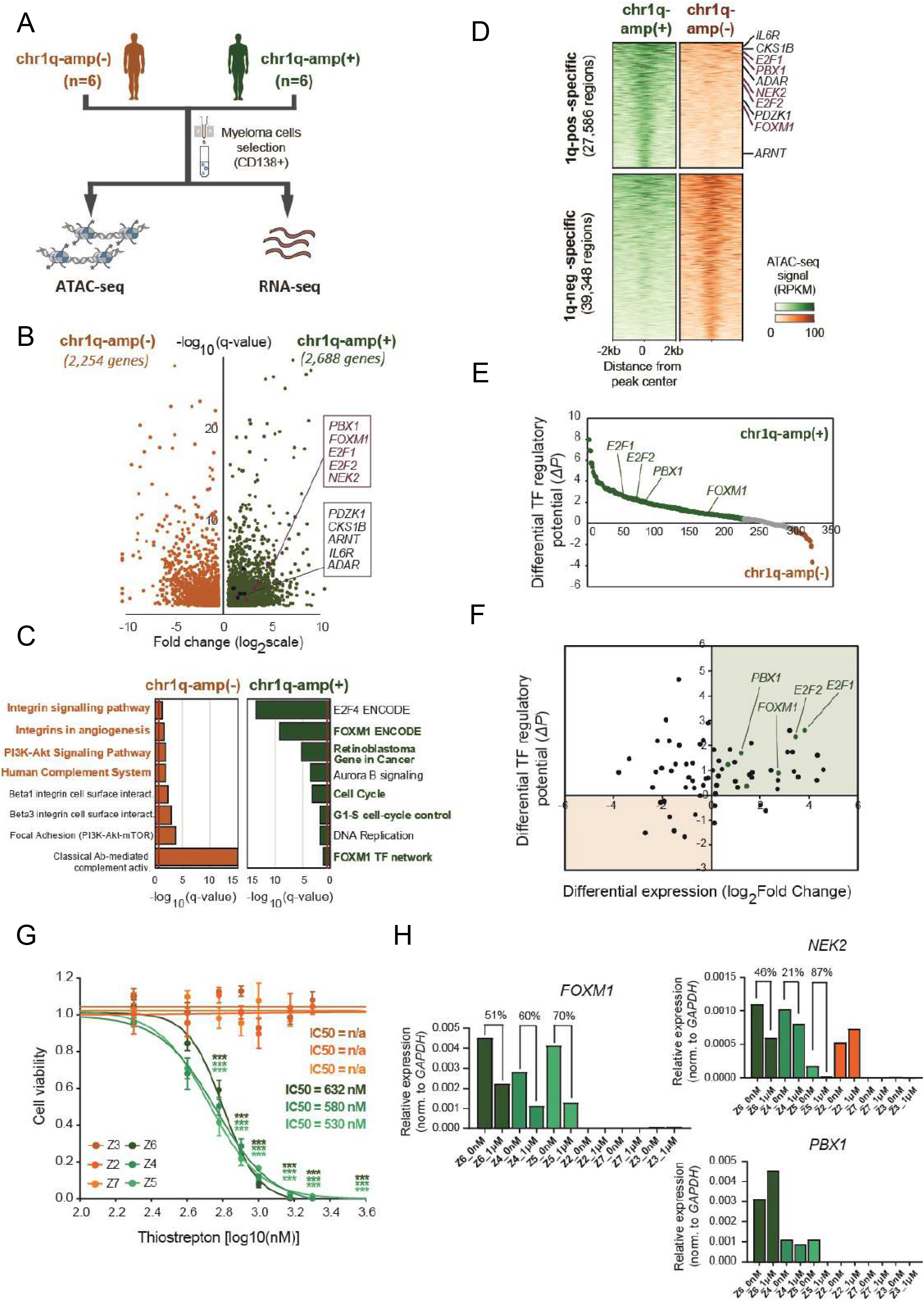
Differential regulome and thiostrepton cytotoxicity profiling of primary chr1q-amplified versus non-amplified MM cells. **(A)** Schematic representation of experimental strategy. Myeloma plasma cells were isolated via magnetic beads selection (CD138+) from bone marrow biopsy samples derived from 6 chr1q-amplified (chr1q-amp(+)) and 6 non-amplified (chr1q-amp(-) MM patients. Differential regulome (TF expression and wiring) analysis was performed via parallel chromatin accessibility (ATACseq) and transcriptome (RNA-seq) profiling. **(B)** Volcano plot displaying differentially expressed genes (chr1q-amp(+), green; chr1q-amp(-), orange). Genes implicated in chr1q-amp pathogenesis in this study (pink) or previous studies (black) are indicated here. **(C)** Enrichment analysis (NCI-Nature pathways) of differentially expressed genes in two patient subgroups. **(D)** Differential ATAC-seq analysis between chr1q-amp(+) and chr1q-amp(-) myeloma plasma cells. Increased accessibility was found on genetic loci of prominent genes upon chr1q amplification (as indicated here). **(E)** Differential ATAC-seq footprinting analysis of expressed TFs in chr1q-amp(+) versus chr1q-amp(-) cells (*ΔP*: differential regulatory potential). TFs of interest are indicated here. **(F)** Scatter plot representation of differential expression (x-axis) and differential regulatory potential (y-axis) of 63 TFs displaying significant differences in both dimensions. Green quartile: TFs with increased expression and *ΔP* in chr1q-amp(+) cells; orange quartile: TFs with decreased expression and *ΔP* in chr1q-amp(+) cells. Key transcription factors are also highlighted here. **(G)** Selective sensitivity of chr1q-amp(+) (n=3, green) versus chr1q-amp(-) (n=3, orange) primary myeloma plasma cells to thiostrepton at 48h after treatment. IC_50_ values were calculated for each patient sample using a non-linear fitting model (fitting line represented here). ****, *P<0.0001.* **(H)** Transcriptional profiling (RT-qPCR) of *FOXM1* and *NEK2* mRNA levels in chr1q-amp(+) (green) and chr1q-amp(-) (orange) primary samples 24h upon thiostretpon (1μM) or mock (0nM) treatment. The (%) decrease in *FOXM1* and *NEK2* mRNA levels is also indicated here.

In addition, we validated functional activation of the PBX1 and shared PBX1-FOXM1 regulatory programmes in *PBX1*-amplified MM cells in a large cohort of patients (MMRF, n=813) and confirmed significant co-expression of *PBX1* and *FOXM1* with almost all of their gene targets across patients in two different cohorts (MMRF, Arkansas; **Supplementary Fig. S6A)**. Importantly, the majority of genes previously shown to comprise high-risk disease signatures in MM (13, 26, 28, 29) were found to be directly regulated by PBX1 (**Supplementary Fig. S6B**). Together, these findings strongly support the critical role of PBX1-FOXM1 axis in promoting proliferative regulatory circuitries determining adverse prognosis and high-risk disease in chr1q-amp MM patients.

### Targeted therapy against chr1q-amp in cancer using a novel, selective PBX1 inhibitor

As the PBX1-FOXM1 axis acts as a central regulatory hub for chr1q-amp MM cells, we next sought to explore the prognostic impact and therapeutic potential of selective PBX1 targeting in chr1-amp cells across several types of cancer. For this purpose, we first analysed transcriptomic data from multiple patient cohorts and found that activation of the PBX1-dependent regulatory signature predicts adverse prognosis in multiple myeloma and 12 solid tumour patient cohorts, including breast, ovarian, lung and brain cancer, in which chr1q-amp is a frequent CNA (**Fig. 6A and Supplementary Fig. S7A**). Next, we tested the impact of our novel, recently reported small-molecule drug T417, which specifically inhibits PBX1 binding to its cognate DNA motif(30), on chr1q-amp cancer cells. We screened four myeloma (MM1.S, U266, NCU.MM1, OPM2), two breast (MCF-7, LTED), two ovarian (OVCAR3, A2780), two lung (A549, H69AR) and one brain (SNB-75) cancer cell lines harbouring at least one additional chr1q copy (**Supplementary Fig. S7B**). Cell viability assays revealed sensitivity of all cell lines to T417 at low μM concentrations (4-28μM), while no significant toxicity was detected upon treatment with the inactive analogue/pro-drug compound DHP52 in two myeloma and two ovarian cancer cell lines (**Fig. 6B and Supplementary Fig. S7C**). In addition, cell cycle analysis revealed significant depletion of the G2/M phase along with G0/1 phase arrest upon T417 treatment (**Fig. 6C**). RT-qPCR-assessed mRNA levels of the PBX1-regulated *FOXM1, NEK2* and *E2F2* genes showed their significant decrease upon treatment with T417 in almost all 11 cell lines (**Fig. 6D**). Interestingly, a significant decrease of *PBX1* mRNA itself was also detected in 8 out of 11 cell lines. This, in conjunction with the binding of PBX1 to its own promoter and putative enhancer, are consistent with a potential mechanism of *PBX1* transcriptional autoregulation (**Fig. 4B**) which would potentiate activity of T417 activity in chr1q-amp cells. Next, using a subcutaneous xenograft myeloma model, we also validated the anti-tumoral activity of T417 *in vivo.* We observed significantly reduced tumour size and weight in the T417-treated versus control mice, while in explanted myeloma cells we detected cell cycle arrest and mRNA depletion of the PBX1-regulated genes (**Fig. 6E-6G** and **Supplementary Fig. S7D-S7H**). In addition, selective cytotoxicity of T417 was detected against *PBX1*-expressing primary chr1q-amplified myeloma cells (X1-X3; n=3), but not against non-amplified MM (X4,X5; n=2) or normal donor peripheral blood B cells (PBBC; n=1) with undetectable *PBX1* mRNA levels (**Fig. 6G and 6H and Supplementary Fig. S7I and S7J**).

**Fig. 6.**
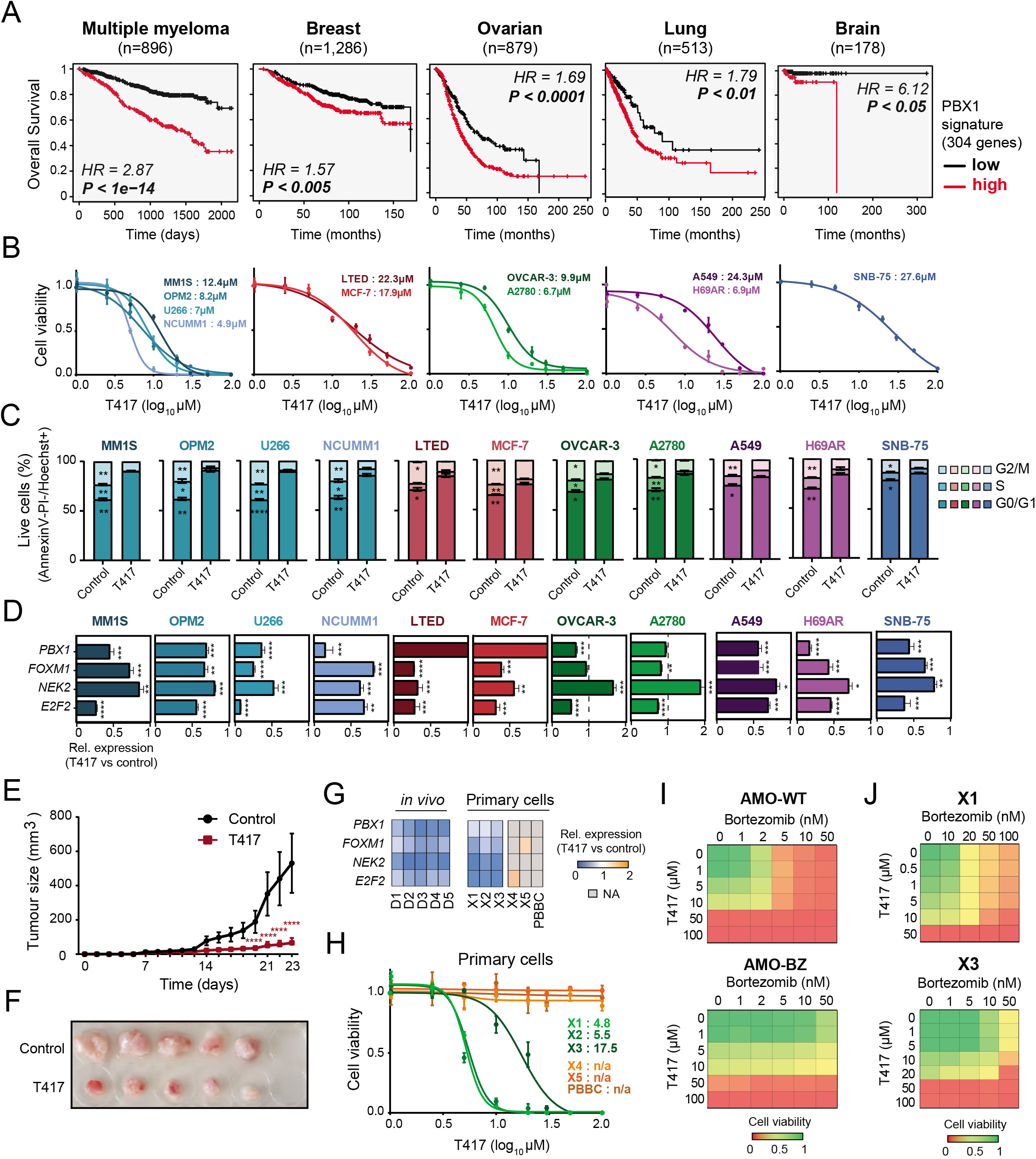
Selective targeting of chr1q-amplified tumour cells with a selective PBX1 inhibitor. **(A)** Survival analysis of multiple myeloma, breast, ovarian lung and brain cancer patient cohorts based on the PBX1 signature expression (red: high, black: low). Kaplan–Meier plots and statistical analysis depict the significantly poor survival outcome of patients with active PBX1 signature. **(B)** Cytotoxicity profiles (n=3) of multiple myeloma (MM1.S, OPM2, U266, NCU.MM1), breast (MCF-7, LTED), ovarian (OVCAR-3, A2780), lung (A549, H69AR) and brain (SNB-75) cancer cell lines 48h after treatment with a selective, small-molecule PBX1 inhibitor (T417). Three independent experiments were performed per cell line and IC_50_ values were calculated using a non-linear fitting model (fitting line represented here). **(C)** Cell cycle profiling of 11 cancer cell lines 48h after treatment with 1% DMSO (control) or T417 (20μM). Three independent experiments were performed per cell line. Asterisks indicate statistical comparisons performed using a two-way non-parametric ANOVA with post-hoc multiple comparisons test. **(D)** Assessment of *PBX1, FOXM1, NEK2* and *E2F2* mRNA levels in 11 cancer cell lines 16-20h after treatment with 1% DMSO (control) or T417 (20μM). Bar graphs illustrate transcriptional levels normalized to corresponding control samples (n=3 replicates). Analysis performed using paired t-test. **(E)** Tumour volumes (mm^3^) of the MM1.S xenografts measured in vehicle- (control) and T417-treated (10 mg/kg/injection) mice across experimental timepoints. Statistical analysis was performed using a two-way ANOVA with post-hoc multiple comparisons test. **(F)** Photograph of tumour sizes dissected at termination date (Day23) from control- and T417-treated mice. **(G)** Heatmap representation of PBX1, FOXM1, NEK2 and E2F2 mRNA levels assessed by RT-qPCR. For the *in vivo* experiment in tumour explanted cells, values represent pairwise comparisons of T417 group (D1-D5) against vehicle-treated group (C1-C5). For *in vitro* primary myeloma plasma cell samples, values represent T417-treated (20μM) versus control-treated (1% DMSO) cells. Grey values correspond to non-applicable (NA) comparisons due to undetectable mRNA levels in control-treated cells. **(H)** Cell viability of primary chr1q-amplified MM (X1,X2,X3; green), non-amplified MM (X4,X5; orange) and normal donor peripheral blood B cells (PBBC; orange) at 48h after treatment with 1% DMSO (control) or T417 (20μM). Non-linear fitting and IC_50_ calculations were performed as described in **(B)**. **(I & J)** Combined cytotoxicity profiling of T417 with Bortezomib in parental (AMO.1-WT), bortezomib-resistant MM cell lines (AMO.1-BZ), and primary chr1q-amp MM cells (X1, X3). Heatmaps represent the cell viability relative to control (1% DMSO) cells. **(K)** Schematic diagram of the overall strategy followed in this study: Construction of an integrated multi-omics and clinical data platform identified 103 genes as candidate pathogenic drivers with prognostic impact in chr1q amplification. Regulatory genomics, genetic and pharmacological approaches revealed a PBX1-FOXM1 axis regulating oncogenic circuitries that promote the proliferative phenotype and high-risk nature of chr1q-amplification in cancer. Selective inhibition of PBX1-FOXM1 axis with existing (thiostrepton) or new (T417) pharmacological agents reveals the translational insights and therapeutic potentials for CNA-targeted therapies in cancer.

Finally, we investigated the potential benefits of T417 treatment combination with proteasome inhibitors, which form the backbone of numerous widely-used regimens to treat newly diagnosed and relapsed multiple myeloma. Combined T417-Bortezomib sensitivity assay as assessed by cell viability performed in parental (AMO.1-WT) and Bortezomib-resistant (AMO.1-BZ) cells showed restoration of AMO.1-BZ cell sensitivity to bortezomib in the presence of T417; for example, combination of 50nM Bortezomib with 10μM T417 in AMO.1-BZ is equivalent or better than 2nM and 10μM respectively in the parental cell line (**Fig. 6I and 6J and Supplementary Fig. S7K-S7M**). The benefit of the dual treatment was also confirmed in two primary chr1q-amp MM samples with known clinical resistance to bortezomib (X1,X3), exemplifying the direct translational applications of T417 in clinic. Overall, these findings highlight the clinical potential of T417 against chr1q-amplified cancer cells as an adjuvant approach against high-risk, chemotherapy-resistant tumours.

## Discussion

Recurrent, high frequency CNA such as chr1q-amp are major oncogenic drivers shared across different types of cancer (1–3). However, delineating the prognostic and functional role of hundreds to thousands of genes and downstream oncogenic pathways associated with specific CNA for development of targeted therapies remains an unmet challenge.

In this study, we focused on chr1q-amp, the most frequent CNA linked to high-risk MM (4–7).

First, by combining WGS and 3D genome data we found that genetic amplification disrupts a large proportion of the chromatin structure throughout the chr1q arm. This level of disruption likely reflects contribution of multiple mechanisms to structural changes in chr1q (31), including isochromosome formation (32), hypoxia-driven tandem duplications (33), jumping translocations (5), chromothripsis and chromoplexy (34), and combination of the above (35). Nevertheless, we detected four main blocks of co-amplification (hyper-domains) which are the product of distinct amplification patterns and retain their overall chromatin structure across MM patients. Of those, only two hyper-domains (B1, B4) contribute to adverse prognosis, and therefore have potential implications in the chr1q-amp biology.

In contrast to previous studies which traditionally focused on 1q21 band alone (9–13), here we employed a large-scale, integrative analysis of clinical and multi-omics datasets (genomics, 3D-genome, epigenomics, transcriptomics and clinical variables) to identify adverse prognosis driver genes across the whole chr1q arm. This analysis validated previously reported high-risk markers in 1q21 locus (9–12), but also linked novel genes to adverse prognosis and highlighted the biological and prognostic significance of two other new areas, the 1q22 and 1q23.3 bands. Collectively, the adverse prognosis genes identified across chr1q are predicted to promote cell cycle and proliferation, suggesting their direct involvement in the well-characterized proliferative phenotype associated with chr1q-amp in MM (13, 28).

Identification of *PBX1,* located in the 1q23.3 region, as a novel candidate driver of high-risk prognosis in chr1-amp MM, also exemplifies the potential of our approach for biological discovery. Indeed, the role of PBX1 in promoting cancer cell survival, metastasis and drug resistance has been reported (20–22, 36). Here we found ectopic expression of PBX1 associated with genetic amplification and strong epigenetic activation of its entire TAD (including proximal and distal DNA elements), suggesting a selective process acting on a whole regulatory domain rather than the gene alone, as previously suggested in oncogenesis (37). Moreover, our composite genetic, epigenetic and pharmacological approaches establish the mechanisms and regulatory networks through which PBX1 regulates the activity of FOXM1, a master TF promoting cell cycle progression (25, 27). The proliferative circuitries regulated by PBX1 and the PBX1-FOXM1 axis are of wider importance in cancer, as they exert a powerful prognostic impact in several cancers. Pertinently, chr1-amp is one of the most frequent CNA not only in MM but also other cancers, including breast and ovarian cancer (20–22).

The finding that pharmacological abrogation of the PBX1-FOXM1 axis selectively impacts survival of chr1q-amp myeloma cells is one of the most notable findings of this work. As well as providing proof-of-principle for developing CNA-specific therapeutic approaches, our data strongly support the central role of PBX1 and FOXM1 in regulation of the transcriptional programme driving the proliferative phenotype and adverse prognosis in chr1q-amp MM. In addition, these findings support our recent efforts for development of T417, a small-molecule inhibitor of PBX1 binding to its cognate DNA motif (30) and suggest the potential benefit of its use in MM and other cancers with chr1-amp. Indeed, along with the previously reported pre-clinical activity of T417 against ovarian cancer (30), our data demonstrate selective targeting against MM, breast, lung, liver and brain cancer cells with chr1-amp. These findings not only validate the presence of a common, PBX1-FOXM1 axis underlying chr1q-amp that is active in many types of cancer, but also provide the basis for clinical development of T417 as a chr1q-amp-targeting therapy.

In summary, we showed that our systems medicine dissection of CNA in cancer, which includes integration of genetic, epigenetic, transcriptional and 3D-chromatin profiles, is a powerful strategy for discovery of genes and cellular oncogenic pathways of biological significance and clinical impact (**Fig.6k**). Through this process, we show that the ectopically expressed PBX1, in co-operation with FOXM1, is a critical driver of the proliferative phenotype in chr1q-amp MM and several other cancers, and we provide proof-of-principle for selective therapeutic targeting of chr1q-amp, the most prevalent CNA in cancer.

## Methods

### Cell cultures

The cell lines MM.1S, U266, NCU.MM1, OPM2, AMO.1-WT, AMO.1-BZ, OVCAR3, A2780, H69AR, NSB-75 and peripheral blood B cells (PBBC) were cultured in RPMI+10% FBS (Gibco), supplemented with 2mM L-glutamine (Sigma), 500IU/mL penicillin and 500μg/mL streptomycin (Sigma), X1 non-essential amino acids (Sigma) and 1mM sodium pyruvate (Sigma), in 37 °C at 5% CO_2_. The same medium with addition of 10ng/ml IL-6 (Gibco, Ref: PHC0066) was used for primary MM plasma cell cultures. MCF-7 cells were cultured in DMEM (Gibco) supplemented with 10% FBS and 500IU/mL penicillin and 500μg/mL streptomycin (Sigma). LTED cells were cultured in phenol-red free DMEM supplemented with 10% FBS and 500IU/mL penicillin and 500μg/mL streptomycin (Sigma). HEK293T cells were cultured in DMEM (Sigma) + 10% FBS (Gibco), supplemented with 2mM L-glutamine (Sigma), 500IU/mL penicillin and 500μg/mL streptomycin (Sigma).

### Primary samples

Bone marrow aspirate samples from multiple myeloma patients and peripheral blood sample from normal donor were obtained upon a written informed consent and under research ethics committee approval (Research Ethics Committee Reference: 11/H0308/9). Bone marrow aspirates were subjected to red cell lysis. Multiple myeloma plasma cells were purified after two rounds of CD138 immunomagnetic selection (Miltenyi Biotech) following the manufacturer’s instructions. Pre- and post-selection purity was assessed by flow-cytometric analysis (BD LSR-Fortessa) using a panel of fluorochrome-labelled anti-CD138, -CD45, -CD19, -CD56 and -CD38 monoclonal antibodies. Purified cells were immediately processed for ATAC-seq and RNA-seq analysis or stored in FBS + 10% DMSO at −150°C for later use.

Mononuclear cells from normal donor peripheral blood sample were isolated by Ficoll-Hypaque (Sigma-Aldrich) density centrifugation following the manufacturer’s instructions. The mononuclear cell interphase layer was aspirated, washed with 1ml PBS, centrifuged at 300g for 5min and resuspended in 100μl PBS. Peripheral blood B cells (PBBC) were isolated using the human Total B cell isolation kit II (Miltenyi Biotec) as per manufacturer’s instructions.

### Molecular cloning

A modified pLKO.1 lentiviral vector (Addgene plasmid #27994), in which the puromycin marker gene was replaced by *eGFP* (for knockdown experiments) or *eBFP* (for rescue experiments) genes. All shRNA oligos we cloned, as previously described (38): scrambled (scrbl) control, 5’-CCTAAGGTTAAGTCGCCCTCG-3’; P11 (anti-*PBX1*), 5’-CGAAGCAATCAGCAAACACAA-3’; P31 (anti-*PBX1*), 5’-ATGATCCTGCGTTCCCGATTT-3’;O1 (anti-*FOXM1*), 5’-CTCTTCTCCCTCAGATATAGA-3’;O4 (anti-*FOXM1*), 5’-GCCAATCGTTCTCTGACAGAA-3’. Successful cloning of recombined vectors was initially confirmed via diagnostic PCR, using the DreamTaq Green PCR Master Mix (2X) (Thermo Scientific) protocol and the 5’-TGGACTATCATATGCTTACCGTAAC-3’ (F) and 5’-GTATGTCTGTTGCTATTATGTCTA-3’ (R) primers, followed by 1% agarose gel electrophoresis. The DNA sequence of positive clones was further confirmed via Sanger Sequencing (outsourced to GeneWiz Ltd), using the same primers set.

The MIGR1-eGFP retroviral vector (Addgene) was used for overexpression experiments. The *PBX1* cDNA sequence (Ensembl, ENSG00000185630) was modified by introducing silent mutations at the shRNA-targeting sites (**Supplementary Methods**) and by adding flanking EcoRI (5’-end) and XhoI (3’-end) restriction enzyme sites for cloning purposes. The designed nucleotide sequence was synthesized and cloned in a pUC57 vector by GenScript Biotech. Both MIGR1 and pUC57-PBX1 vectors were digested using EcoRI and XhoI (FastDigest, Thermo Scientific), purified by gel extraction and ligated in a molarity ratio of 1:2 (vector:insert). Ligation mixtures were transformed into *E. Coli* competent cells and amplified using the GeneJET Plasmid Maxiprep Kit (Thermo Scientific). Diagnostic PCR with the MIGR1 primers set, followed by 1% agarose gel electrophoresis, was performed to obtain positive recombinant clones and their exact sequence was confirmed via Sanger sequencing (GeneWiz Ltd) using four different primers: (F1) 5’-CCTAAGCCTCCGCCTCCTCTTC-3’ ; (R1) 5’-GAAGACAGGGCCAGGTTTCCGG-3’; (F2) 5’-TTAGATCTCTCGAGATGGACGAGCAGCCCAG-3’ ; (R2) 5’-GGGCGGAATTCTCAGTTGGAGGTATCAGAGTGAAC-3’.

### Virus production

Recombinant lentiviral and retroviral vectors produced from previous steps were amplified using the GeneJET Plasmid Maxiprep Kit (Thermo Scientific). The 3^rd^ generation lentiviral (pRSV.REV, pMDLgpRRE, pMD2.VSVG, Addgene) and 2^nd^ generation retroviral (pUMVC3-gag-pol, pMD2.G-VSVG, Addgene) helper plasmids were also amplified using the same kit. The pLKO.1 vectors were cotransfected with lentiviral helper plasmids into HEK293T cells using the calcium phosphate transfection method (39). For retrovirus production, MIGR-EV (original MIGR1 construct) or MIGR-*PBX1* vectors were co-transfected with retroviral helper plasmids, following the same protocol. Medium was removed after 8h and cells were treated with 10ml glycerol (15% v/v) for 3min, washed with PBS and incubated in fresh medium. Viral supernatant was collected and concentrated at 48- and 72-hours post-transfection via ultracentrifugation at 23,000 rpm for 2h at 4°C. Viral pellets were resuspended in FBS-free DMEM medium overnight at 4°C under constant shaking. For long-term storage, virus was aliquoted into separate tubes, immediately incubated for 15min in dry ice and stored at −80°C.

### Cell transduction experiments

For knockdown experiments, MM1.S and U266 myeloma cells were transduced with shRNA-*eGFP*-containing pLKO.1 lentiviruses in 24-well plates (10 x 10^4^ cells per construct) and in presence of polybrene (Sigma; final concentration 8μg/ml). Medium was changed by centrifugation (5min, 300xg) 16h post-transduction and cell viability was monitored 48h later (Day3) and every 48-72h on the basis of GFP expression using the BD LSR FORTESSA flow-cytometry analyser. To determine the knockdown efficiency, transduced cells were purified 3 days post-transduction on the basis of GFP expression by fluorescence activated cell sorting (FACS) using a BD FACS AriaIII sorter (MRC flow-cytometry facility, Imperial College London). Total RNA extraction and transcriptomic analysis of isolated cells performed as detailed below. For cell cycle analysis, transduced cells were collected 6 days (for PBX1) or 5 days (for FOXM1) post-transduction and subsequently processed according to the protocol below.

For overexpression experiments, MM.1S and NCU.MM1 cells were transduced with MIGR1-EV or MIGR1-*PBX1* retrovirus with the addition of polybrene (Sigma; final concentration 8μg/ml); cell medium was changed by centrifugation (5min, 300xg) 20h post-infection. Long-term cell viability was assessed via trypan-blue staining and flow-cytometry (based on cell viability and GFP intensity) using the BD LSR FORTESSA analyser. Transcriptomic analysis of transduced cells was performed as detailed below.

For rescue experiments, MM.1S cells already containing the MIGR1-EV and MIGR1-*PBX1* vectors were transduced with shRNA-*eBFP*-containing lentiviruses as described above; cells were isolated 3 days post-transduction based on dual GFP/BFP markers fluorescence.

### Cell cycle analysis

Cell cycle analysis was performed on live cells as previously described (38). For knockdown, *in vivo* and T417 cytotoxicity experiments, transduced cells were cultured at 37 °C for 60min in the presence of the Hoechst 33342 live-cell staining dye (Abcam, USA) to a final concentration of 10μM. Next, cells were collected via centrifugation (5min, 300xg), washed twice with 1x Annexin V buffer (eBiosciences, USA) and incubated with Annexin V antibody (eBiosciences, USA) for 15min at 4oC in the dark. Finally, Propidium iodide (Sigma, USA) was added (final concentration 250μg/ml) in cell mixture and flowcytometric analysis was performed using the BD LSR FORTESSA. For phenotype rescue experiments, the Vybrant™ DyeCycle™ Ruby live-cell staining dye (final concentration 10μM, Thermo Scientific) was used along with Propidium iodide.

### Intracellular staining

PBX1 intracellular staining for analysis by flow cytometry was performed as described (40), with minor modifications. All incubations were performed on ice and shielded from light. After harvesting, cells were washed and resuspended in 100μl PBS, fixed by adding equal volume of 4% formaldehyde solution (16% methanol-free formaldehyde by Polysciences, 18814, diluted in PBS) whilst vortexing to ensure single cell suspension, and incubated for 3h. Cells were then span at 600xg for 5min and washed twice with PBS. At the final wash, care was taken to remove all supernatant. Cell permeabilization was performed by adding 100μL of stain buffer and incubating for 30min (stain buffer: 5% BSA and 0.5% Triton X-100 (Sigma-Aldrich) in PBS). Subsequently cells were stained with 1μg of primary antibody, either anti-PBX1 or isotype control, and incubating on ice for 45min in the dark. After another wash with stain buffer (600xg, 5min), cells were resuspended in 100μL and stained with 2.5μL of secondary antibody, incubating for 45min. Cells were finally washed twice with stain buffer, resuspended in 300μl PBS and analysed on a BD LSR FORTESSA analyser. Upon analysis, the Median Fluorescence Intensity (MFI) ratio was identified for each sample, denoting ratio of median fluorescence of anti-PBX1 antibody over isotype control. Antibodies used: anti-PBX1 (Abnova, clone 4A2, H00005087-M01), isotype control mouse IgG2a k (eBioscience 14-4724-81), secondary antibody APC-conjugated rat anti-mouse IgG2a (eBioscience 17-4210-80).

### Drug sensitivity and cytotoxicity assays

Cancer cell lines or primary cells were plated in 96-well plate at a density of 3×10^3^ cells/well in triplicate, and treated with 0nM (control) or various concentrations of thiostrepton (B7336-APE-50mg, Stratech); or, 1% DMSO (control) or various concentrations of T417 inhibitor (provided by Dr. Wang) as indicated and cultured for 48h. Myeloma cells (AMO1-WT, AMO1-BZ, primary MM cells) were treated with 1% DMSO (control) or various combinations of T417 and Bortezomib (Cell Signaling Technology) concentrations and cultured for 24h. Cell viability was tested by a CellTiter-Glo assay (Promega) using a microplate reader (Fluostar, BMG, Durham, NC). Drug cytotoxicity curves were obtained from non-linear fit analysis in GraphPad Prism and IC_50_ was defined as the concentration that results in a 50% decrease in the number of live cells. Total RNA extraction and RT-qPCR (described below) were performed 16-20h after treatment with 0nM (control) or 1μM thiostrepton; or, 1% DMSO (control) or 20μM T417. Cell cycle analysis (described above) was performed 24h after treatment with 1% DMSO (control) or 20μM T417.

### *in vivo* experiments

For the *PBX1* knockdown experiment, nine female and nine male NOD.Cg-Prkdcscid Il2rgtm1Wjl/SzJ (NSG), 8-10 week-old mice were purchased from Charles River UK Ltd. Maintenance and experiments were performed at Imperial College London Animal Facility, in accordance with the 1986 Animal Scientific Procedures Act and under a United Kingdom Government Home Office-approved project license (PPL/PP8553679). Human MM1.S myeloma cells were transduced with pLKO.1-scrbl, pLKO.1-P11 or pLKO.1-P31 lentiviral vectors (as previously described) and collected two days posttransduction (~6×10^6^ cells per construct) in 1ml PBS, washed twice with PBS after centrifugations at 300xg and resuspended in 300ul PBS. 1×10^6^ cells (corresponding to 50ul) per construct were aliquoted into Eppendorf tubes and mixed along with 100μl of Matrigel Basement membrane LDEV-free matrix (Scientific laboratory supplies) on ice. Cells were resuspended gently within the mixtures and injected subcutaneously into the mice, in such a way that each construct was transplanted in 3 male and 3 female mice. Monitoring after injections was performed every 48h and body weights were measured using an analytical scale. Tumour growth was observed and measured using a caliper ruler 2-3 times per week by using the formula: Tumour volume = (length x width^2^) / 2 (length represents the longest diameter and width represents the perpendicular diameter of the tumour). Experiment was terminated when tumours reached the maximum allowed size (≤15mm in length or width). Upon termination, all mice were culled on the same day, tumours were immediately dissected and photographed. Tumour sizes were measured post-mortem with the use of caliper and tumour weights measurements were obtained using an analytical scale. Finally, tumours were homogenized using a plunger of a 1ml syringe and filtered through a 40μm cell strainer (Cole-Parmer) twice, to isolate single cells. Approximately 10% of the cells obtained from each tumour sample were stained with antihuman HLA-ABC-APC (Miltenyi: 130-101-466) mAb and analysed using the BD FORTESSA flow cytometer.

For the PBX1 inhibitor experiment, six male and six female NSG mice were purchased from Charles River UK Ltd. Approximately 10×10^6^ cells per mouse were resuspended in PBS, mixed with Matrigel Basement membrane LDEV-free matrix (Scientific laboratory supplies) as described above and injected subcutaneously into the mice. After daily monitoring, all tumours reached a measurable size 7 days post-injection and mice were randomized to include 3 male and 3 female mice per treatment group. Mice were treated via intraperitoneal route with Control (vehicle): 1% DMSO (Sigma), 10% 1-methyl-2-pyrrolidone (Sigma), 40% polyethylene glycol 400 (PEG; Sigma), 50% PBS; or PBX1 inhibitor: T417 (10mg/kg/injection) + vehicle, following an intermittent schedule of 4 days on / 3 days off per week for a total of 10 treatments. Tumour sizes and mouse body weights were monitored daily, as described above. When tumours reached the maximum allowed size, experiment was terminated and all mice were culled on that day. Tumour dissections, size and weight measurements and cell homogenizations were performed as mentioned above. Approximately 40% of tumour cells were stained with the anti-human HLA-ABC-GFP (Miltenyi: 130-101-466) mAb; two-thirds were FACS-sorted for total RNA extraction and one-third was plated in 24-well plates (1ml RMPI +10% FBS +1% PS + 1% NEA +1% SP) and subjected to cell cycle analysis, as previously described.

### Immunohistochemistry (IHC)

Immunohistochemical analysis of trephine biopsy samples from multiple myeloma patients and tonsil tissues from healthy donors was performed by the Histopathology unit of Hammersmith Hospital. In short, serial 4μm sections from formalin fixed, paraffin embedded human trephine and tonsil tissues were sliced on to Superfrost Plus® slides (VWR) and incubated at 60°C for 45min. Slides were dewaxed by immersion in Histo-Clear (National Diagnostics USA) and rehydrated with subsequent immersion in 100% ethanol, 70% ethanol and distilled water. Antigen retrieval was performed by immersion in 95°C TRIS-EDTA (Sigma-Aldrich Ltd) comprised of 1M Tris-HCl (pH approximately 8.0) containing 0.1M EDTA for 30min in a Grant SUB Aqua 5 Plus water-bath. Slides were rinsed in PBS and endogenous peroxidase activity blocked using 0.3% hydrogen peroxide (Sigma-Aldrich Ltd) in PBS for 15min. Thereafter, slides were rinsed with PBS and incubated with 1.5% normal goat serum (Vector Laboratories) for 30min prior to incubation with primary antibody PBX1 clone 4A2 (Abnova, 1mg/mL; 1/50), at room temperature for one hour. Slides were rinsed in PBS and incubated with secondary biotinylated antibody (Goat Anti-Mouse IgG, Vector Laboratories, 1/100) for 30min followed by an avidin/biotin peroxidase complex (VECTASTAIN Elite ABC Kit, Vector Laboratories) for 30min Chromogenic reaction was developed using DAB (Diaminobenzidine, Vector ImmPACT DAB Peroxidase Substrate) for 3min then halted by immersion in running tap water for 5min. Nuclei were counterstained with Gill 2 Haematoxylin (Thermo-Scientific Shandon) and blued in Scott’s tap water (in-house preparation) for 1min. Slides were dehydrated in 70% ethanol and 100% ethanol, cleared in Histo-Clear (National Diagnostics USA) and mounted in DPX (VWR BDH ProLab). Slides were allowed to dry in the fume hood for about 30min; microscopic examination and high-resolution photography were performed at the Histopathology laboratory in Hammersmith Hospital.

### Chromatin Immunoprecipitation (ChIP)-qPCR and ChIP-seq

Chromatin Immunoprecipitation was performed as previously described with minor modifications (41). The antibodies used in this study include: monoclonal anti-PBX1 (4A2 clone, M01, Abnova), monoclonal anti-FOXM1 (ab1: GTX102170, GeneTex; ab2: Gtx100276_c3, GeneTex), monoclonal anti-IgG (clone 3E8, Santa Cruz). In brief, MM1.S or U266 cells were crosslinked with 1% formaldehyde (Sigma-Aldrich) for 15min. Glycine (1.25M) was used to quench the formaldehyde. Cells were washed with PBS and lysed in ChIP lysis buffer (40mM Tris-HCl (pH:8), 4mM EDTA (pH:8), 1% (v/v) Triton-X 100, 300mM NaCl supplemented with 1x protease inhibitors (Thermoscientific)) for 10min on ice. Cell lysates were sonicated on a Diagenode BioRuptor sonicator (No of Cycles: 40; Intensity: High, 30sec on/ 30sec off). The size of the sheared chromatin was assessed by agarose gel electrophoresis after reverse crosslinking to an average length of 300-500bp. Input genomic DNA and IP DNA was prepared by treating aliquots of chromatin with RNase (Thermoscientific), proteinase K (NEB) and heat for decrosslinking followed by Ampure XP beads (Beckman Coulter) purification.

Quantitative real-time PCR for ChIP assays (ChIP-qPCR) was performed using the SYBR Green Master Mix (Thermo Scientific) along with target-specific primers in optical 96-well plates on ABI StepOne Plus Thermocycler (Applied Biosystems) with the following settings: 50°C (2min), 95°C (2min), 95°C (3sec) and 60°C (30sec) alternating for 45 cycles, 95°C (15sec), 60°C (1min) and 95°C (15sec). Primers linearity and specificity was determined before use. Relative enrichment over input was calculated using the 2-ΔCt method and values were compared with corresponding IgG controls. Primers sets: PBX1-promoter, (F) 5’-ACCGTCTGTGTTCTTTCGTGT-3’, (R) 5’-CTTTCCCTGCTCGCCTTACT-3’; NEK2-promoter, (F) 5’-ATCTCGCAGTCTATTGGCAGG-3’, (R) 5’-GGTTAAAAGCAGACGCCGAC-3’; NEK2-enhancer (F) 5’-CACCACCACCATCTTTGCAC-3’, (R) 5’-ACACGTTATGTCCTCTGGGC-3’; FOXM1-enhancer (F) 5’-TCATTCACCGGTTGATGCCT-3’, (F) 5’-GTGGTTGTTGGTGGAACAGC-3’.

For high-throughput sequencing experiments, ChIP and input DNA libraries were prepared for amplification by using the NEBNext Ultra II ChIP-seq Library Prep Master Mix Set for Illumina (NEB) following manufacturer’s protocols with no modifications. The quantity was determined using the Qubit High Sensitivity DNA kit (Life Technologies) and library size was determined using the Bioanalyser High Sensitivity DNA kit (Agilent). Finally, libraries were quantified using the Universal Library Quantification Kit for Illumina (Kapa Biosystems) and run on the ABI StepOne Plus Thermocycler (Applied Biosystems).

Sequencing was performed at the Genomics Facility at MRC LIMS of Imperial College London using the Illumina HiSeq 2500 platform to obtain single-end 50bp reads.

### Reverse transcription and RT-qPCR

Total RNA was isolated from cell lines using the Nucleospin RNA kit (Macherey-Nagel) and quantified using NanoDrop 1000 (Thermo Fisher Scientific, Appendix 2). Synthesis of cDNA was achieved using RevertAid First Strand cDNA Synthesis Kit (Thermo Scientific) and quantitative real-time PCR was performed using the Taqman Real-Time Assays reagent (Thermo Fisher) in Fast optical 96-well plates on an ABI StepOne Plus Thermocycler (Applied Biosystems) as follows: 50°C (2min), 95°C (10min), 95°C (15sec) and 60°C (1min) alternating for 45 cycles. Transcription levels were evaluated with the comparative threshold cycle (Ct) method and following the 2-ΔΔCt method with normalization to *GAPDH* housekeeping gene expression. The taqman probes (Thermo Scientific) used in this study are: *PBX1* (Hs00231228_m1), *E2F2* (Hs01007097_m1), *FOXM1* (Hs01073586_m1), *NEK2* (Hs00601227_1), *GAPDH* (Hs03929097_g1).

### RNA-seq

Total RNA was extracted from FACS-sorted myeloma cells using the Nucleospin RNA kit (Macherey-Nagel). The Qubit RNA Assay kit (Life Technologies) was used to determine the RNA quantity. Quality of RNA extracts was assessed on the Bioanalyser using the RNA pico kit (Agilent). Samples with RIN value higher than 8 were processed using the NEBNext poly(A) mRNA Magnetic Isolation kit and the NEBNext Ultra II RNA Library Prep kit for Illumina (New England Biolabs), following manufacturer’s instructions. The Qubit High Sensitivity DNA kit (Life Technologies) was used for libraries quantification; library size was evaluated using the Bioanalyser High Sensitivity DNA kit (Agilent). Libraries from the same experiment were diluted to 5nM, pooled together and sequenced at the BRC Genomics Facility (Imperial College London) using the Illumina HiSeq 4000 platform to obtain paired-end 75bp reads.

### ATAC-seq

ATAC-seq was performed as previously described (42). Briefly, 50,000 purified plasma cells, myeloma plasma cells or cell lines, were washed with cold PBS (Sigma) at 500g at 4°C for 5 min. The cells were resuspended in 50 μL of cold Lysis Buffer (10 mM Tris-HCl, pH 7.4, 10mM NaCl, 3 mM MgCl2, 0.1% IGEPAL CA-630) and washed at 500g at 4°C for 10min. The nuclei were subjected to transposase reaction for 30min at 37°C; termination of the reaction and DNA purification was performed using a MiniElute Kit (Qiagen) and eluted twice with 10 μL. The purified DNA was amplified as described before with NEBNext High-Fidelity 2x PCR Master Mix (New England Biolabs). The PCR amplified product was cleaned twice with (0.9X) AMPure beads (Beckman). The quality of the libraries was assessed with the Bioanalyzer High Sensitivity DNA kit (Agilent). The libraries were quantified using the NEBNext Library Quant Kit for Illumina (New England Biolabs) on a StepOne Plus Real-Time PCR (Applied Biosystems). The libraries were sequenced at the Genomics Facility at ICL using the Illumina HiSeq 4000 platform to obtain paired-end 75bp reads.

### Bioinformatics and clinical informatics analysis

All methods used for bioinformatics and clinical informatics analyses are described in the **Supplementary methods** file.

### Statistical analysis and additional software

Statistical analyses for all biological experiments was performed using GraphPad Prism software, with the appropriate test applied for each experiment. Flow cytometry and FACS data acquisition was done using the BD FACSDiva™ software, and analysis was later performed using the FlowJo X software. For cloning strategies design and *in silico* evaluation of DNA sequences, SnapGene (GSL Biotech LLC) was used when necessary.

## Data and code availability

High-throughput sequencing data generated during this study have been deposited to the Gene Expression Omnibus repository (GEO): MMCL ChIP-seq and RNA-seq files (GSE165060) and primary MM ATAC-seq files (GSE153381).

Code used in this study can be accessed from the specified github page: https://github.com/nikostrasan/PBX1-project

## Authors’ Disclosures

The authors declare no conflict of interest.

## Authors’ Contributions

NT designed study, conceived and implemented computational pipelines, designed and performed experiments, wrote manuscript. AlK, KP, YS, IK, BB, and KK performed experiments. XX assisted in bioinformatics data analysis. PD and NK performed and interpreted IHC analysis. RMS assisted in clinical informatics data analysis. AC and HWA provided clinical samples. IAGR, TL and LM wrote manuscript. VSC and AnK: designed study, supervised experiments, generated draft manuscript. All authors contributed to the final draft of the manuscript.

## Acknowledgements

NT, VC, XX and KP were supported by Bloodwise (Blood Cancer UK), AlK was supported by Kay Kendall Leukaemia Fund and Imperial NIHR Biomedical Research Centre. We also acknowledge support from LMS/NIHR Flow-cytometry Facility, Imperial NIHR Biomedical Research Centre Genomics Facility, Imperial NIHR Biomedical Research Centre, Cancer Research UK Experimental Cancer Medicine Centre. The NCU.MM1 myeloma cell line was kindly provided by Dr. Ichiro Hanamura (Aichi Medical University, Japan). The OVCAR3 and A2780 cell lines were kindly provided by Dr. Sadaf Ghaem-Maghami (Imperial College London, UK). We would also like to thank Dr. Udayakumar Achandira (Hammersmith hospital) for his kind advice and assistance with the MMCL FISH analysis.

## Notes

### Competing Interest Statement

The authors have declared no competing interest.

