## Supplementary Figure for "Systems medicine dissection of chromosome 1q amplification reveals oncogenic regulatory circuits and informs targeted therapy in cancer"

### 3. Supplemental figures

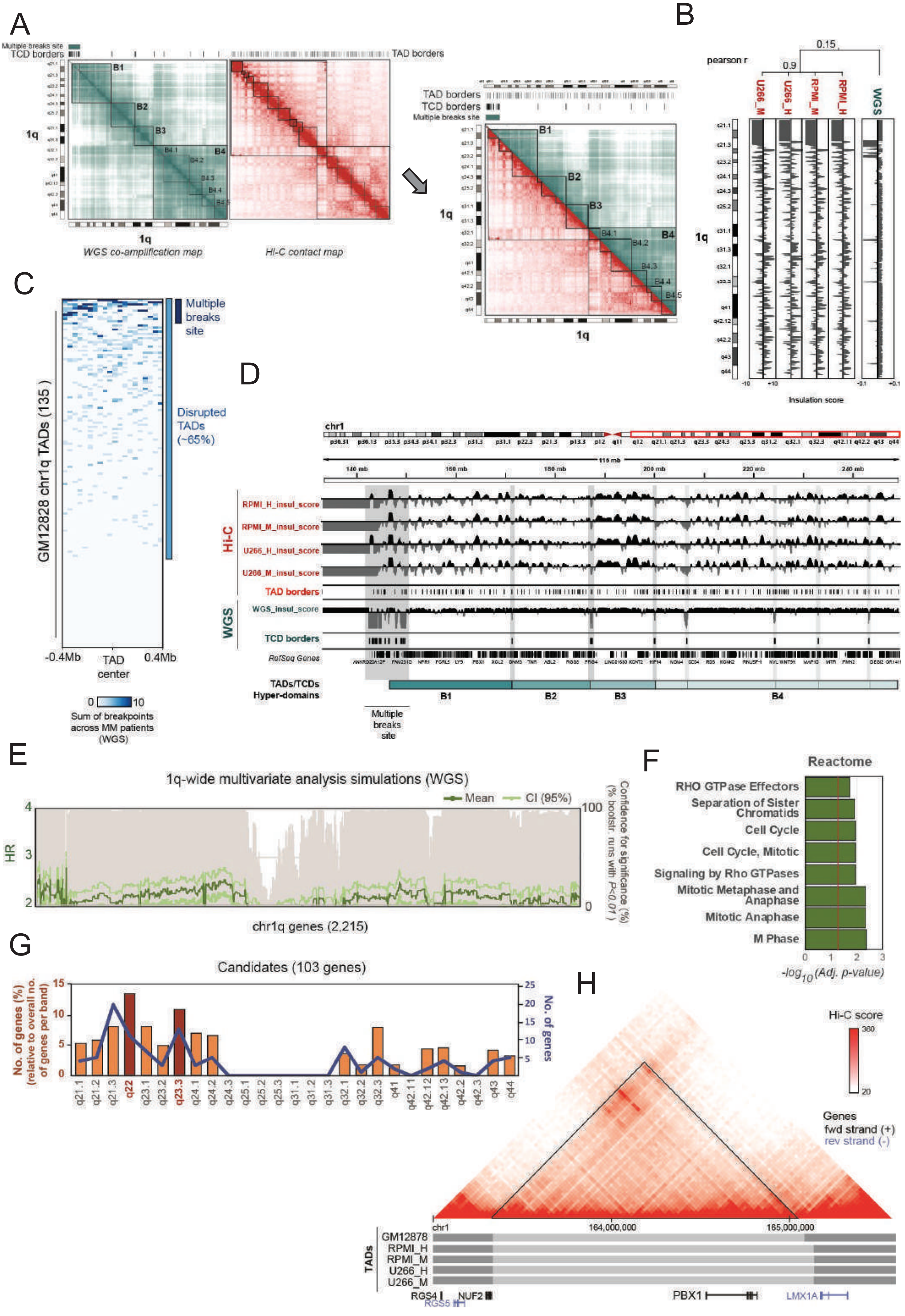

A
B
C
D
E
F
G
H
Multiple breaks site
TCD borders
TAD borders
WGS co-amplification map
Hi-C contact map
pearson r
U266_M
U266_H
RPMI_M
RPMI_H
WGS
Insulation score
Multiple breaks site
Disrupted TADs (~65%)
GM12828 chr1q TADs (135)
-0.4Mb
TAD center
0.4Mb
Sum of breakpoints across MM patients (WGS)
RPMI_H_insul_score
RPMI_M_insul_score
U266_H_insul_score
U266_M_insul_score
TAD borders
WGS_insul_score
TCD borders
RefSeq Genes
TADs/TCDs Hyper-domains
1q-wide multivariate analysis simulations (WGS)
Mean
CI (95%)
HR
Confidence for significance (% bootstr. runs with P<0.01)
chr1q genes (2,215)
Reactome
RHO GTPase Effectors
Separation of Sister Chromatids
Cell Cycle
Cell Cycle, Mitotic
Signaling by Rho GTPases
Mitotic Metaphase and Anaphase
Mitotic Anaphase
M Phase
-log10(Adj. p-value)
Candidates (103 genes)
No. of genes (%) (relative to overall no. of genes per band)
No. of genes
Hi-C score
Genes
fwd strand (+)
rev strand (-)
GM12878
RPMI_H
RPMI_M
U266_H
U266_M
TADs
RGS4
RGS5
NUF2
PBX1
LMX1A

**Supplementary Fig. S1 related to Fig. 1. Supplementary information on systems medicine analysis of chr1q amplification in myeloma.**

**(A)** WGS co-amplification (cyan; 2D genome) and HiC contact (red; 3D genome) maps (top) and merged 2D-3D map (bottom) of chr1q in multiple myeloma cells. TCDs and TADs are indicated by the vertical bars on top of each map, respectively. Vertical bars on top of maps indicate TCD and TAD borders. The multiple breakpoints locus (pericentromeric area with genetic instability) is also indicated with a blue bar. Four conserved hyperdomains (B1-4) were identified after overlaying the two maps.

**(B)** Insulation score profiles of Hi-C (red) and WGS co-amplification (cyan) data across the chr1q arm. Cladogram indicates the correlation (pearson r) between experiments.

**(C)** Heatmap illustrates the total number of WGS amplification breakpoints in MM patients (MMRF dataset) with reference to non-amplified B-cell TADs (GM12828; [Wu et al, 2017].

**(D)** IGV snapshot of chr1q locus displays an overview of the insulation scores and TAD/TCD borders identified across chr1q in this study. Red, 3D genome Hi-C data analysis (obtained from Wu et al., 2017); cyan, WGS co-amplification data analysis (MMRF dataset). RPMI_H: RPMI8226 HindIII, RPMI_M: RPMI8226 MobI, U266_H: U266 HindIII, U266_M: U266 MobI.

**(E)** Multivariate survival analysis overview of all chr1q genes across 859 MMRF patients against 73 genetic markers. The mean (forest-green) and 95% CI (light-green) of Hazard Ratio (HR) estimations upon Monte-Carlo simulations (5,000 iterations) per gene are represented here. Grey bars represent the percentage (%) of bootstrapping tests returning significant (*P<0.01*) survival risk for each gene.

**(F)** Reactome pathway analysis of all 103 candidate driver genes.

**(G)** Distribution of candidate genes (103) across cytogenetic bands of chr1q arm. Line graph (blue) represents the absolute number of genes; bar graph (orange) indicates the number of genes per band (relative to their gene density). Bars in red indicate two bands with the highest relative enrichment for candidate driver genes.

**(H)** Hi-C interaction map (top) and TAD (bottom) of GM12878 B lymphoblastoid cell line at the genetic area of *PBX1*. Preservation of the *PBX1* TAD is shown as compared to TAD from MM cell lines (U266, RPMI8226)

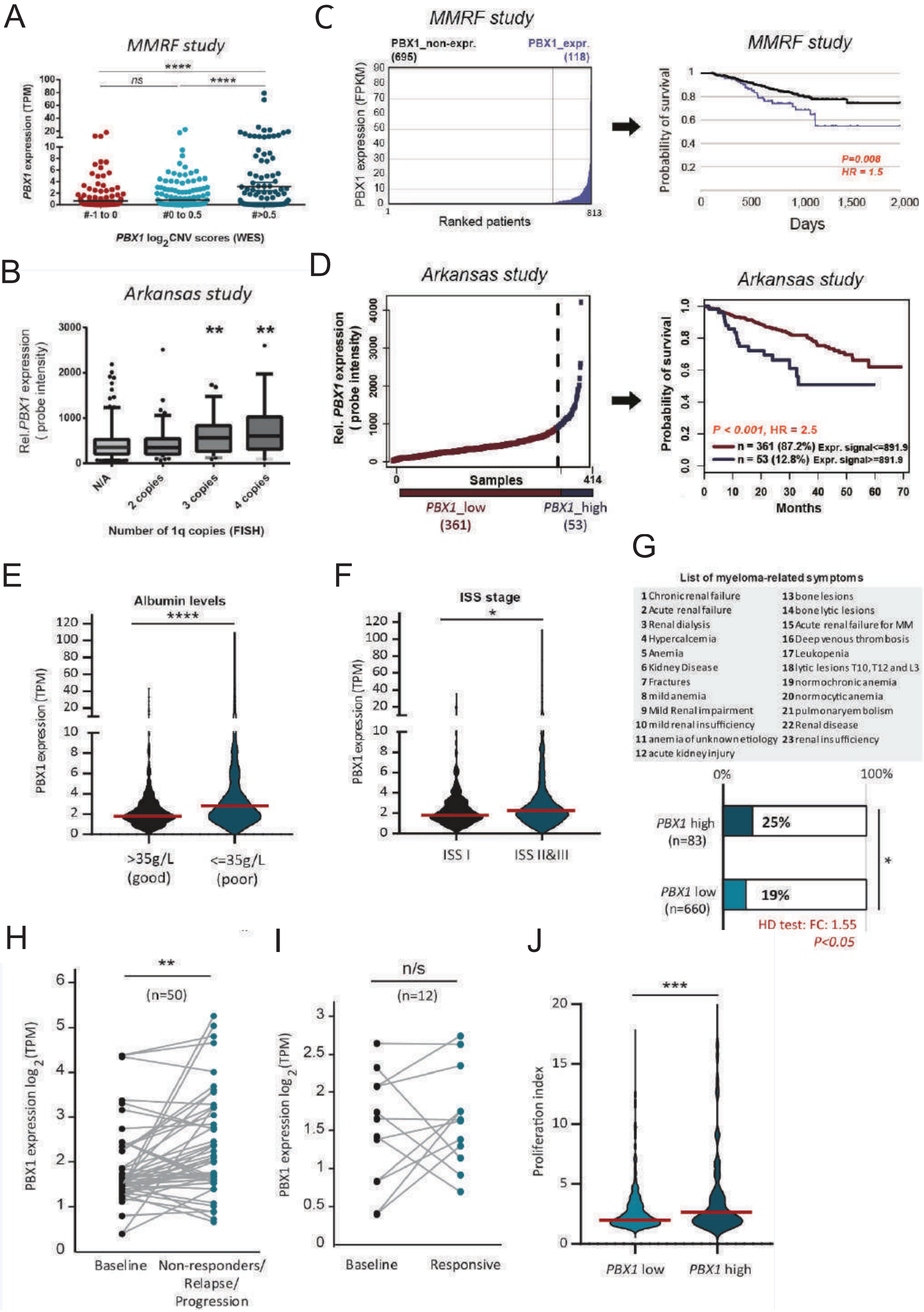
A
MMRF study
****
ns
****
PBX1 expression (TPM)
#-1 to 0
#0 to 0.5
#>0.5
PBX1 log2CNV scores (WES)
B
Arkansas study
Rel.PBX1 expression ( probe intensity)
**
**
N/A
2 copies
3 copies
4 copies
Number of 1q copies (FISH)
C
MMRF study
PBX1_non-expr. (695)
PBX1_expr. (118)
PBX1 expression (FPKM)
Ranked patients
813
MMRF study
Probability of survival
P=0.008
HR = 1.5
Days
D
Arkansas study
Rel. PBX1 expression ( probe intensity)
Samples
414
PBX1_low (361)
PBX1_high (53)
Arkansas study
Probability of survival
P < 0.001, HR = 2.5
n = 361 (87.2%) Expr. signal<=891.9
n = 53 (12.8%) Expr. signal>=891.9
Months
E
Albumin levels
****
PBX1 expression (TPM)
>35g/L (good)
<=35g/L (poor)
F
ISS stage
*
PBX1 expression (TPM)
ISS I
ISS II&III
G
List of myeloma-related symptoms
1 Chronic renal failure
2 Acute renal failure
3 Renal dialysis
4 Hypercalcemia
5 Anemia
6 Kidney Disease
7 Fractures
8 mild anemia
9 Mild Renal impairment
10 mild renal insufficiency
11 anemia of unknown etiology
12 acute kidney injury
13 bone lesions
14 bone lytic lesions
15 Acute renal failure for MM
16 Deep venous thrombosis
17 Leukopenia
18 lytic lesions T10, T12 and L3
19 normochronic anemia
20 normocytic anemia
21 pulmonary embolism
22 Renal disease
23 renal insufficiency
0%
100%
PBX1 high (n=83)
25%
PBX1 low (n=660)
19%
*
HD test: FC: 1.55
P<0.05
H
**
(n=50)
PBX1 expression log2(TPM)
Baseline
Non-responders/ Relapse/ Progression
I
n/s
(n=12)
PBX1 expression log2(TPM)
Baseline
Responsive
J
***
Proliferation index
PBX1 low
PBX1 high

**Supplementary Fig. S2. Related to Fig. 1. *PBX1* as a high-risk prognostic biomarker in MM.**

**(A)** Genetic amplification of *PBX1* is associated with its overexpression in MM patients. WES and RNA-seq data obtained from MMRF study and patients were stratified according to their *PBX1* copy numbers (CNV log2 ratio: -1 to 0; 0 to 0.5; >0.5).

**(B)** Relative PBX1 probe intensities from DNA microarray expression data from the Arkansas study are displayed for each patient, based on the number of 1q copies detected by FISH analysis.

**(C)** MMRF patients stratified according to PBX1 expression (RNA-seq) into high and low expressing groups; survival analysis between PBX1 high vs low patient groups (HR: Hazard ratio)

**(D)** DNA microarray expression data from Arkansas study used to stratify MM patients according to their relative PBX1 probe intensity (blue:PBX1 high, red:PBX1 low); subsequent Kaplan-Meier plot and survival analysis of the two cohorts shows significant differences in disease outcome. Log-rank test was applied for survival analyses.

(**E & F**) Violin plots illustrating the *PBX1* levels of MM patients stratified based on (e) albumin levels and (f) the International Staging System (ISS) stage**.** Classification: albumin >35g/L: good prognosis, <=35g/L: poor prognosis; ISSI: good prognosis, ISSII&III: poor prognosis.

**(G)** Overrepresentation analysis of clinical symptoms in newly diagnosed patients at presentation with respect to their *PBX1* expression levels; box with 23 myeloma-related symptoms (top) and bar graphs (bottom) displaying the percentage of patients with high or low PBX1 expression with at least one of the annotated myeloma symptoms.

**(H & I)** Patient-matched RNA-seq levels of *PBX1* in (**H**) non-responders, relapsed or patients with progressive disease and (**I**) responsive patients, in comparison to their baseline RNA-seq expression. Number of patients are indicated in each plot.

**(J)** Proliferation index of MM patients with low or high PBX1 expression levels.

Statistical comparisons for **(A, B)** were performed using Kruskall-Wallis with Dunn's post-hoc multiple comparisons test. Hypergeometric distribution (HD) test was used for **(G)**. Mann-Whitney test was performed in **(E, F, J)** and paired t-test was used in **(H, I)**. Error bars show standard errors of mean *: *P<0.05;* **: *P<0.01;* ***: *P<0.001;* ****: *P<0.0001; n/s: not significant.*

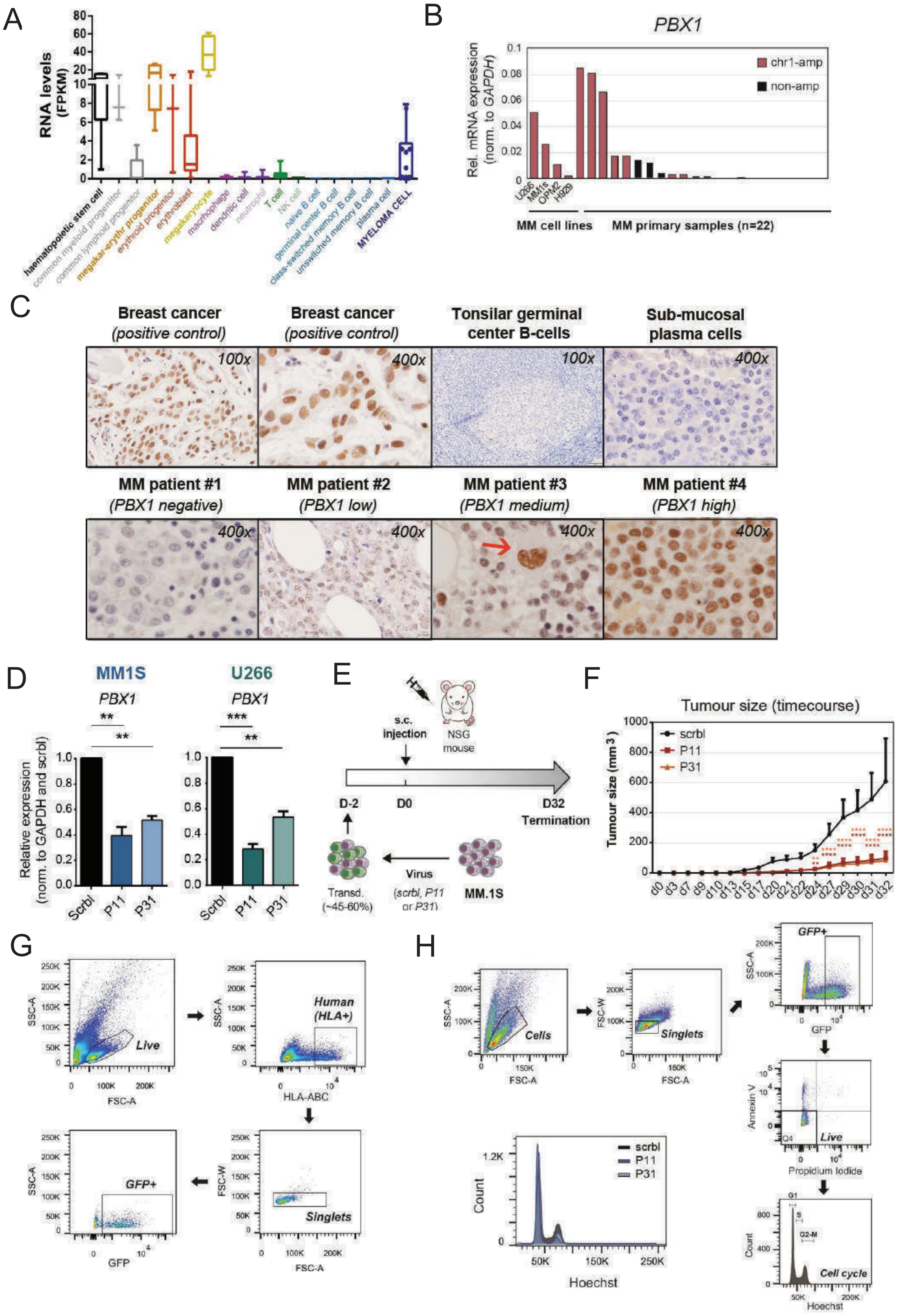

A
RNA levels (FPKM)
haematopoietic stem cell
common myeloid progenitor
common lymphoid progenitor
megakar-erythr progenitor
erythroid progenitor
erythroblast
megakaryocyte
macrophage
dendritic cell
neutrophil
T cell
NK cell
naive B cell
germinal center B cell
class-switched memory B cell
unswitched memory B cell
plasma cell
MYELOMA CELL
B
PBX1
Rel. mRNA expression (norm. to GAPDH)
chr1-amp
non-amp
U266
MM1s
OPM2
H929
MM cell lines
MM primary samples (n=22)
C
Breast cancer (positive control)
Tonsilar germinal center B-cells
Sub-mucosal plasma cells
MM patient #1 (PBX1 negative)
MM patient #2 (PBX1 low)
MM patient #3 (PBX1 medium)
MM patient #4 (PBX1 high)
100x
400x
D
MM1S
U266
PBX1
Relative expression (norm. to GAPDH and scrbl)
Scrbl
P11
P31
E
s.c. injection
NSG mouse
D-2
D0
D32 Termination
Transd. (~45-60%)
Virus (scrbl, P11 or P31)
MM.1S
F
Tumour size (timecourse)
Tumour size (mm3)
scrbl
P11
P31
G
SSC-A
FSC-A
Live
HLA-ABC
Human (HLA+)
FSC-W
Singlets
GFP
GFP+
H
Cells
Singlets
GFP+
Annexin V
Propidium Iodide
Live
Count
Hoechst
scrbl
P11
P31
G1
S
G2-M
Cell cycle

**Supplementary Fig. S3 Related to Fig. 2. The functional role of PBX1 in chr1q-amplified myeloma cells.**

**(A)** *PBX1* mRNA expression profile across hematopoietic development and multiple myeloma plasma cells. RNA-seq data (FPKM values) from 17 primary haematopoietic cell populations and multiple myeloma cells were obtained from the Blueprint Consortium Data Portal. Boxplots illustrate median values and SD.

**(B)** PBX1 mRNA expression (qPCR) in 4 chr1q-amplified MMCL and 22 MM samples.

**(C)** Immunohistochemical analysis of PBX1 expression in healthy donor and cancer patient tissues. Top: Luminal sections (100x; 400x) from breast cancer patients used as positive controls for PBX1 staining. Tonsillar germinal centre B-cells and submucosal plasma cells from healthy tissues stain negative for PBX1 protein expression. Bottom: Bone marrow trephine biopsy samples from multiple myeloma patients containing myeloma cells with no, moderate or high PBX1 expression. Red arrow in patient #2 indicates strong PBX1 staining of megakaryocyte cells within the myeloma patients bone marrow, serving as internal positive control for each sample.

**(D)** qPCR analysis of PBX1 mRNA expression upon knockdown with P11 and P31 shRNAs, compared to scrambled (scrbl) control. Data analysis was performed using a one-way ANOVA with post-hoc multiple comparisons test.

**(E)** Schematic overview of the experimental design used in this study. In short, human MM1.S myeloma cells were transduced with anti-*PBX1* shRNAs-containing (P11, P31) or scrambled control lentiviruses (~45-60% transduction levels) and cells were injected subcutaneously in NSG mice (3 males, 3 females, i.e., n=6 per group). Tumour size was monitored every 48h throughout the course of the experiment, until it reached the maximum allowed volumes (Termination Day 32).

**(F)** Calculated tumour sizes ($mm^3$) of P11, P31 and scrambled control mouse groups across different experimental timepoints. Statistical analysis was performed using a two-way ANOVA with post-hoc multiple comparisons test.

**(G)** Gating strategy for flow-cytometric analysis of isolated tumours on termination day. FACS plots illustrate a characteristic example of Live/HLA+/Singlet/GFP+ populations identified in the tumour of a scrambled-control mouse.

**(H)** Gating strategy followed during flow cytometry analysis. In short, P11, P31 and scrambled shRNA-expressing myeloma cells were analysed 6 days after transduction as follows: (i) total cells (excl. debris) ; (ii) singlets (excl. doublets) ; (iii) GFP+ cells (transduced cells); (iv) Annexin V / Propidium Iodide double negative (Q4, excl. early and late apoptotic cells); (v) G1,S,G2-M phases based on Hoechst staining. **:*P<0.01*; ****P<0.001.; ****P<0.0001.*

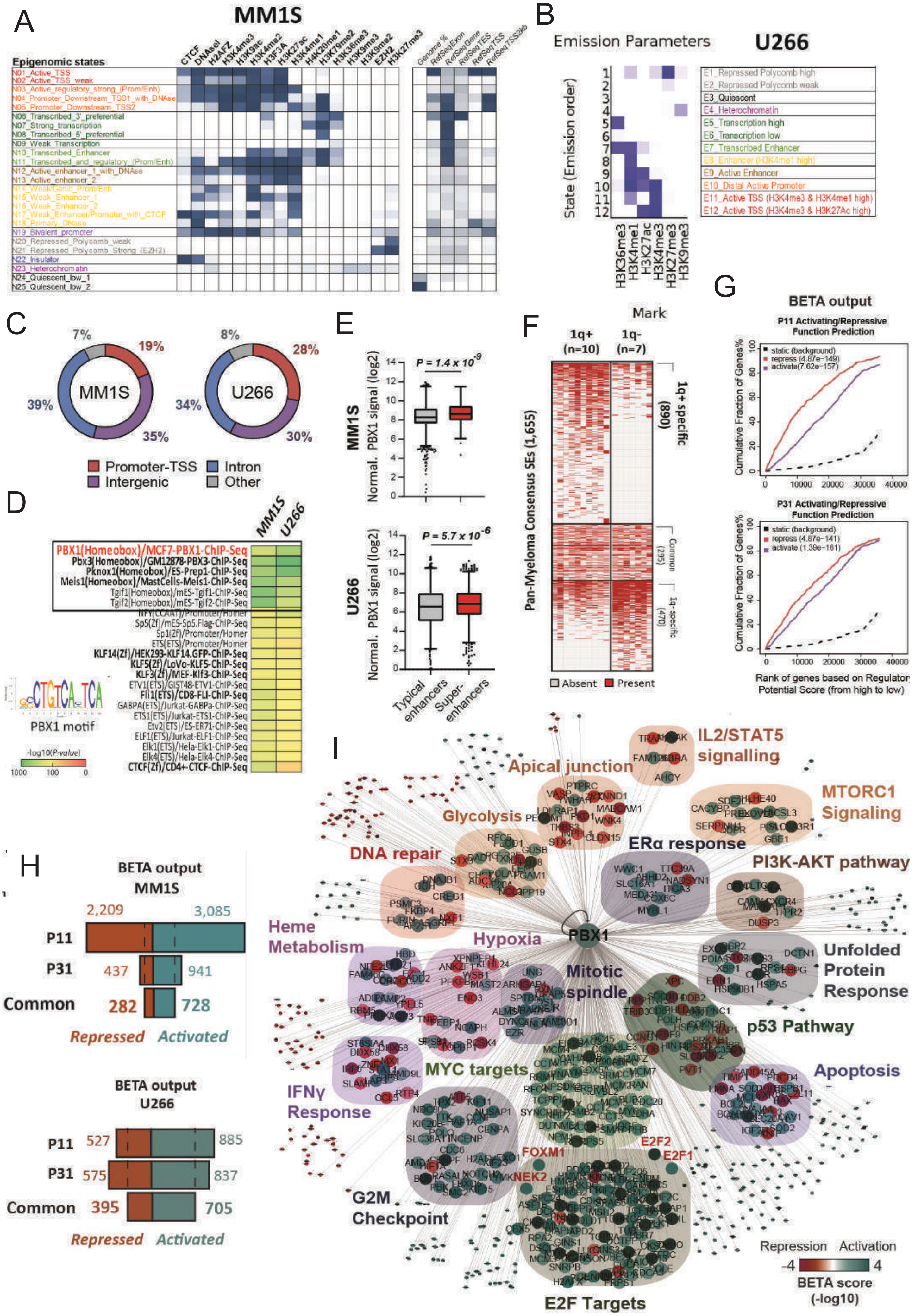
A
MM1S
Epigenomic states
CTCF
DNAseI
H2AFZ
H3K4me3
H3K9ac
H3K4me2
H3F3A
H3K27ac
H3K4me1
H4K20me1
H3K79me2
H3K36me3
H3K9me3
EZH2
H3K9me2
H3K27me3
Genome %
RefSeqExon
RefSeqGene
RefSeqTES
RefSeqTSS
RefSeqTSS2kb
N01_Active_TSS
N02_Active_TSS_weak
N03_Active_regulatory_strong_(Prom/Enh)
N04_Promoter_Downstream_TSS1_with_DNAse
N05_Promoter_Downstream_TSS2
N06_Transcribed_3'_preferential
N07_Strong_transcription
N08_Transcribed_5'_preferential
N09_Weak_Transcription
N10_Transcribed_Enhancer
N11_Transcribed_and_regulatory_(Prom/Enh)
N12_Active_enhancer_1_with_DNAse
N13_Active_enhancer_2
N14_WeakGenic_Prom/Enh
N15_Weak_Enhancer_1
N16_Weak_Enhancer_2
N17_Weak_Enhancer/Promoter_with_CTCF
N18_Primary_DNase
N19_Bivalent_promoter
N20_Repressed_Polycomb_weak
N21_Repressed_Polycomb_Strong_(EZH2)
N22_Insulator
N23_Heterochromatin
N24_Quiescent_low_1
N25_Quiescent_low_2
B
Emission Parameters
U266
State (Emission order)
E1_Repressed Polycomb high
E2_Repressed Polycomb weak
E3_Quiescent
E4_Heterochromatin
E5_Transcription high
E6_Transcription low
E7_Transcribed Enhancer
E8_Enhancer (H3K4me1 high)
E9_Active Enhancer
E10_Distal Active Promoter
E11_Active TSS (H3K4me3 & H3K4me1 high)
E12_Active TSS (H3K4me3 & H3K27Ac high)
H3K36me3
H3K4me1
H3K27ac
H3K4me3
H3K27me3
H3K9me3
C
MM1S
7%
19%
39%
35%
U266
8%
28%
34%
30%
Promoter-TSS
Intron
Intergenic
Other
D
MM1S
U266
PBX1(Homeobox)/MCF7-PBX1-ChIP-Seq
Pbx3(Homeobox)/GM12878-PBX3-ChIP-Seq
Pknox1(Homeobox)/ES-Prep1-ChIP-Seq
Meis1(Homeobox)/MastCells-Meis1-ChIP-Seq
Tgif1(Homeobox)/mES-Tgif1-ChIP-Seq
Tgif2(Homeobox)/mES-Tgif2-ChIP-Seq
NFY(CCAAT)/Promoter/Homer
Sp5(Zf)/mES-Sp5.Flag-ChIP-Seq
Sp1(Zf)/Promoter/Homer
ETS(ETS)/Promoter/Homer
KLF14(Zf)/HEK293-KLF14.GFP-ChIP-Seq
KLF5(Zf)/LoVo-KLF5-ChIP-Seq
KLF3(Zf)/MEF-Klf3-ChIP-Seq
ETV1(ETS)/GIST48-ETV1-ChIP-Seq
Fli1(ETS)/CD8-FLI-ChIP-Seq
GABPA(ETS)/Jurkat-GABPa-ChIP-Seq
ETS1(ETS)/Jurkat-ETS1-ChIP-Seq
Etv2(ETS)/ES-ER71-ChIP-Seq
ELF1(ETS)/Jurkat-ELF1-ChIP-Seq
Elk1(ETS)/Hela-Elk1-ChIP-Seq
Elk4(ETS)/Hela-Elk4-ChIP-Seq
CTCF(Zf)/CD4+-CTCF-ChIP-Seq
PBX1 motif
-log10(P-value)
1000
100
0
E
MM1S
P = 1.4 x 10^-9
Normal. PBX1 signal (log2)
U266
P = 5.7 x 10^-6
Typical enhancers
Super-enhancers
F
Mark
1q+ (n=10)
1q- (n=7)
Pan-Myeloma Consensus SEs (1,655)
1q+ specific (890)
Common (295)
1q- specific (470)
Absent
Present
G
BETA output
P11 Activating/Repressive Function Prediction
static (background)
repress (4.87e-149)
activate(7.62e-157)
P31 Activating/Repressive Function Prediction
static (background)
repress (4.87e-141)
activate (1.39e-161)
Cumulative Fraction of Genes%
Rank of genes based on Regulatory Potential Score (from high to low)
H
BETA output
MM1S
2,209
3,085
P11
P31
437
941
Common
282
728
Repressed
Activated
BETA output
U266
P11
527
885
P31
575
837
Common
395
705
Repressed
Activated
I
PBX1
IL2/STAT5 signalling
Apical junction
MTORC1 Signaling
Glycolysis
DNA repair
ERα response
PI3K-AKT pathway
Heme Metabolism
Hypoxia
Mitotic spindle
Unfolded Protein Response
p53 Pathway
MYC targets
Apoptosis
IFNγ Response
G2M Checkpoint
FOXM1
NEK2
E2F2
E2F1
E2F Targets
Repression
Activation
-4
4
BETA score (-log10)

**Supplementary Fig. S4 related to Fig.3. The epigenetic programme of PBX1 in chr1q-amplified cells.**
**(A & B)** Emission parameters of imputed data illustrate the epigenomic signals derived from 16 **(A)** and 6 **(B)** chromatin marks used to construct ChromHMM maps in MM1.S **(A)** and U266 **(B)** cells.
**(C)** Genomic annotation of PBX1 cistrome in MM1.S (left) and U266 (right) cells.
**(D)** Motif analysis of PBX1 binding sites in MM1.S and U266 cells. The previously known PBX1 transcription factor motif, detected among top hits in this analysis (red), is also depicted here in weblogo representation.
**(E)** Boxplot representation of PBX1 binding signal in typical enhancers (grey) and SEs (red) in MM1.S (top) and U266 (bottom) MM cells.
**(F)** Heatmap representation of distribution of PBX1-bound consensus SEs across 17 MM samples. FISH was used to stratify samples into chr1q-amplified (1q+) and non-amplified (1q-) groups. SEs were clustered based on their presence in the two groups as 1q+ specific, common and 1q-negative specific.
**(G)** Regulatory potential prediction models display significant activating (blue) and repressive (red) function of PBX1 in U266 cells. Models derived from BETA-plus analysis, after integrating the PBX1 ChIP-seq binding sites with P11-depleted (top) and P31-depleted (bottom) RNA-seq analysis.
**(H)** Numbers of genes activated (blue) and repressed (red) directly by PBX1 in MM1.S cells (top) and U266 (bottom) cells
**(I)** The gene regulatory programme of PBX1 in U266 cells. Biological annotation of genes was performed using the Molecular Signatures Database. Node colours represent average predicted activation (blue) or repression (red) for each gene. Transcriptional targets with prominent biological role are highlighted in red font.

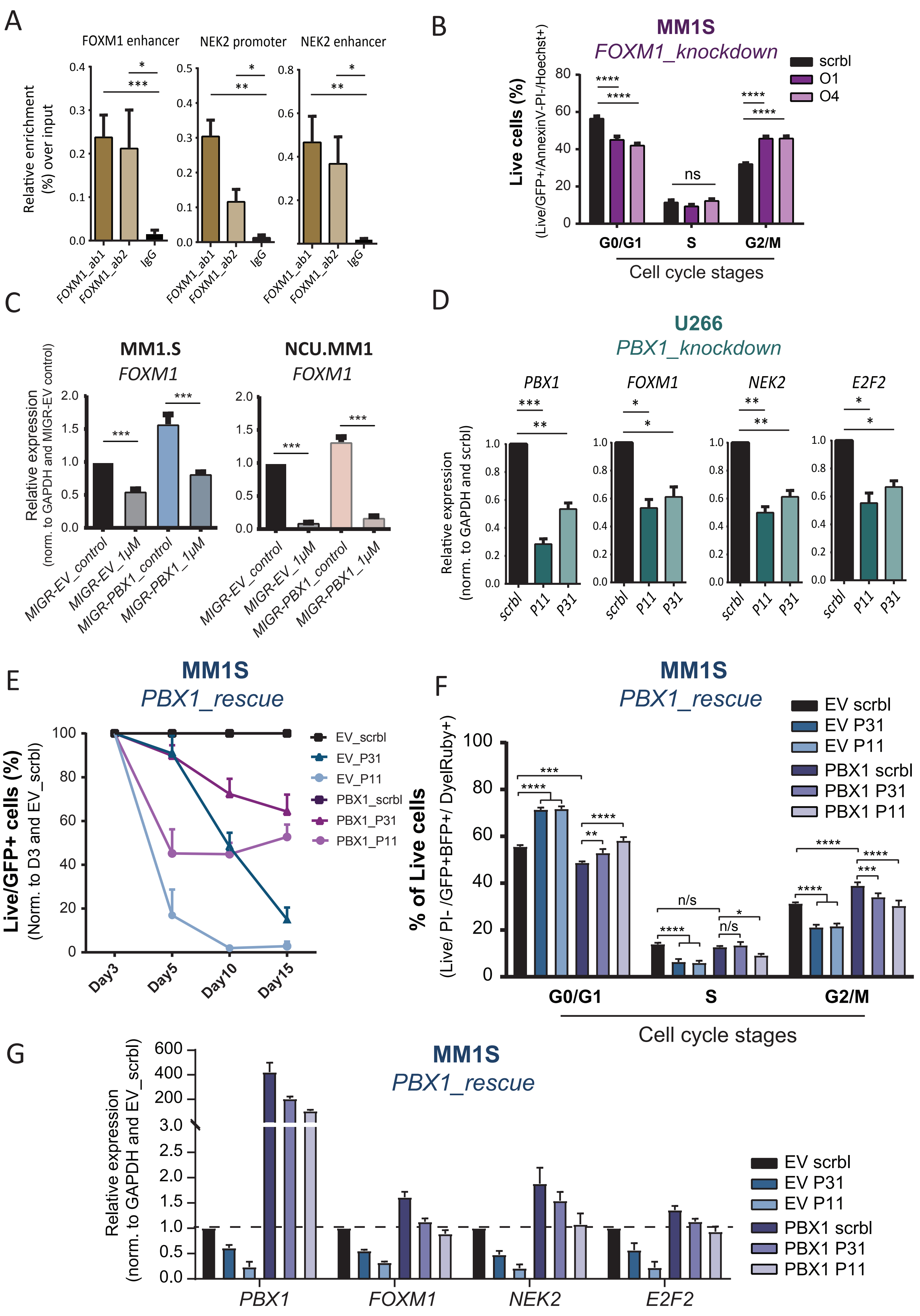
A
FOXM1 enhancer
NEK2 promoter
NEK2 enhancer
Relative enrichment (%) over input
FOXM1_ab1
FOXM1_ab2
IgG
B
MM1S
FOXM1_knockdown
scrbl
O1
O4
Live cells (%)
(Live/GFP+/AnnexinV-PI-/Hoechst+)
G0/G1
S
G2/M
Cell cycle stages
ns
C
MM1.S
NCU.MM1
FOXM1
Relative expression
(norm. to GAPDH and MIGR-EV control)
MIGR-EV_control
MIGR-EV_1μM
MIGR-PBX1_control
MIGR-PBX1_1μM
D
U266
PBX1_knockdown
PBX1
FOXM1
NEK2
E2F2
Relative expression
(norm. to GAPDH and scrbl)
scrbl
P11
P31
E
MM1S
PBX1_rescue
EV_scrbl
EV_P31
EV_P11
PBX1_scrbl
PBX1_P31
PBX1_P11
Live/GFP+ cells (%)
(Norm. to D3 and EV_scrbl)
Day3
Day5
Day10
Day15
F
MM1S
PBX1_rescue
EV scrbl
EV P31
EV P11
PBX1 scrbl
PBX1 P31
PBX1 P11
% of Live cells
(Live/ PI- /GFP+BFP+/ DyelRuby+)
n/s
G0/G1
S
G2/M
Cell cycle stages
G
MM1S
PBX1_rescue
Relative expression
(norm. to GAPDH and EV_scrbl)
EV scrbl
EV P31
EV P11
PBX1 scrbl
PBX1 P31
PBX1 P11
PBX1
FOXM1
NEK2
E2F2

**Supplementary Fig. S5 related to Fig.4. Genetic experiments confirm regulatory hierarchy among *PBX1, FOXM1, NEK2* and *E2F2* in chr1q-amplified MM cells.**

**(A)** ChIP-qPCR analysis against FOXM1 (ab1, ab2 antibodies) and IgG control on prominent DNA elements in MM1.S cells (n=4; the *FOXM1* enhancer, *NEK2* promoter and *NEK2* enhancer regulatory regions as displayed in Fig5b). Statistical analysis was performed using a one-way paired-samples ANOVA with post-hoc multiple comparisons test.

**(B)** Flow cytometry-based, cell cycle analysis of MM1.S cells 5 days after transduction with O1, O4 and scrambled control lentiviruses (n=3). Analysis was performed via one-way ANOVA with post-hoc multiple comparisons test.

**(C)** Relative *FOXM1* mRNA levels (RT-qPCR) in *PBX1*-overexpressing (MIGR-PBX1) or non-overexpressing (MIGR-EV) MM1.S and NCUMM1 cells 24h after treatment with vehicle (control) or 1μM thiostrepton (n=3). Statistical analysis was performed using a pairwise t-tests. Rescue of *PBX1*-depleted phenotype in MM1.S cells.

**(D)** RT-qPCR analysis of *PBX1*, *FOXM1*, *NEK2*, *E2F2* levels upon *PBX1* knockdown in U266 cells. Analysis performed using one-way ANOVA with post-hoc multiple comparisons test.

**(R)** Time-course, flow-cytometric analysis of MM1.S cell survival upon PBX1 overexpression (PBX1) or not (EV) and *PBX1* shRNA-mediated knockdown (P11,P31) or not (scrbl). Data represent the mean values of four independent biological replicates.

**(F)** Cell cycle profiling (n=3) and (**g**) RT-qPCR profiling (n=3) of *PBX1, FOXM1 NEK2* and *E2F2* mRNA expression in control (EV_scrbl), *PBX1*-silenced (EV_P31,EV_P11), *PBX1*-overexpressing (PBX1_scrbl) and *PBX1*-rescued (PBX1_P11, PBX1_P31) cells. Error bars show standard errors of mean *: P<0.05; **: P<0.01; ***: P<0.001; ****: P<0.0001; n/s: not significant.

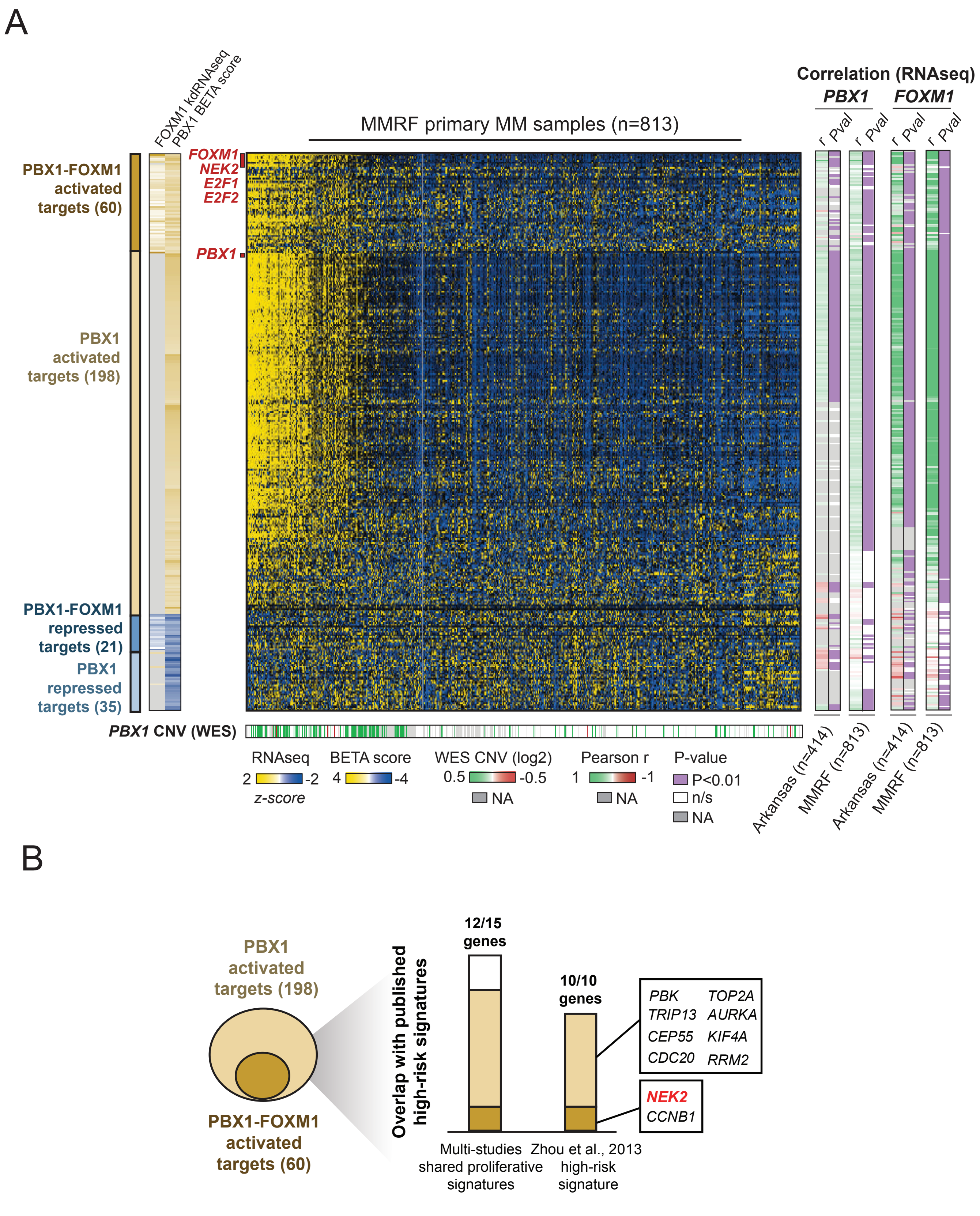
A
FOXM1 kdRNAseq
PBX1 BETA score
MMRF primary MM samples (n=813)
Correlation (RNAseq)
PBX1
FOXM1
r Pval
r Pval
r Pval
r Pval
PBX1-FOXM1 activated targets (60)
PBX1 activated targets (198)
PBX1-FOXM1 repressed targets (21)
PBX1 repressed targets (35)
FOXM1
NEK2
E2F1
E2F2
PBX1
PBX1 CNV (WES)
RNAseq
2 -2
z-score
BETA score
4 -4
WES CNV (log2)
0.5 -0.5
NA
Pearson r
1 -1
NA
P-value
P<0.01
n/s
NA
Arkansas (n=414)
MMRF (n=813)
Arkansas (n=414)
MMRF (n=813)
B
PBX1 activated targets (198)
PBX1-FOXM1 activated targets (60)
Overlap with published high-risk signatures
12/15 genes
10/10 genes
PBK
TOP2A
TRIP13
AURKA
CEP55
KIF4A
CDC20
RRM2
NEK2
CCNB1
Multi-studies shared proliferative signatures
Zhou et al., 2013 high-risk signature

**Supplementary Fig. S6 related to Fig. 5. Activation of the PBX1-FOXM1 regulatory network is linked to high-risk prognosis in multiple myeloma.**

**(A)** Combined epigenomic and transcriptomic analysis reveals the PBX1 (258 predicted activated, 56 repressed target genes) and the PBX1-FOXM1 shared regulatory networks (60 predicted activated, 21 repressed target genes). From left to right: heatmap representations display FOXM1-depleted MM1.S RNA-seq data, average regulatory (BETA) score of PBX1 targets in MM1.S and U266 cells, RNA-seq data of primary MM samples for the MMRF study (n=813), correlation analysis of PBX1 and FOXM1 expression levels against all gene targets with the use of the Arkansas (n=414) and MMRF (n=813) primary MM expression datasets. Bottom: PBX1 Copy Number Variation (CNV) calculated from WES data.

**(B)** Venn diagram illustrates overlap of the identified PBX1 and PBX1-FOXM1 shared gene targets with previously defined proliferative and high-risk myeloma signature genes [23-26]. The 10 ultra-high risk genes from *Zhou et al, 2013* are also shown here.

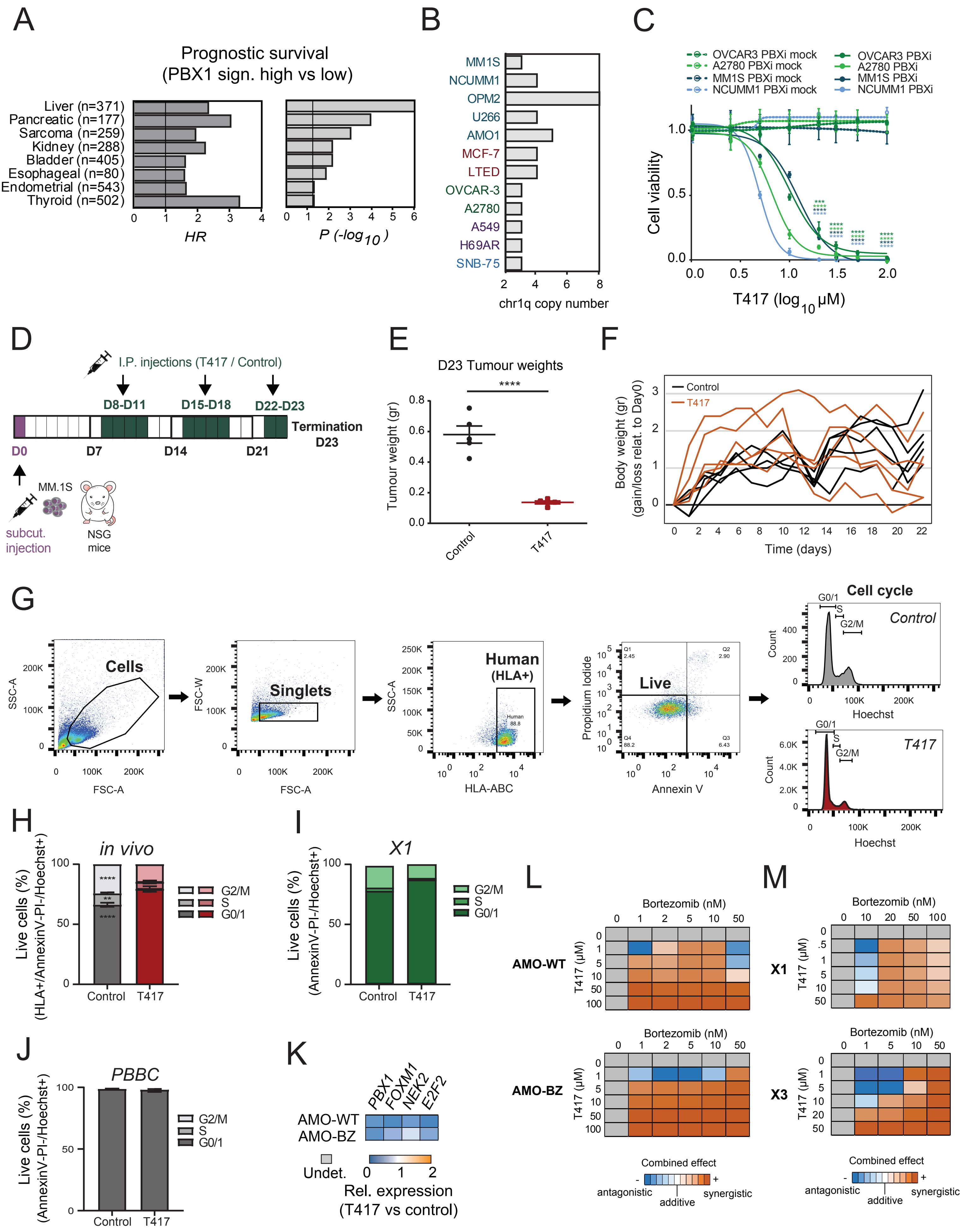
A
Prognostic survival
(PBX1 sign. high vs low)
Liver (n=371)
Pancreatic (n=177)
Sarcoma (n=259)
Kidney (n=288)
Bladder (n=405)
Esophageal (n=80)
Endometrial (n=543)
Thyroid (n=502)
HR
P (-log10)
B
MM1S
NCUMM1
OPM2
U266
AMO1
MCF-7
LTED
OVCAR-3
A2780
A549
H69AR
SNB-75
chr1q copy number
C
OVCAR3 PBXi mock
A2780 PBXi mock
MM1S PBXi mock
NCUMM1 PBXi mock
OVCAR3 PBXi
A2780 PBXi
MM1S PBXi
NCUMM1 PBXi
Cell viability
T417 (log10 μM)
D
I.P. injections (T417 / Control)
D8-D11
D15-D18
D22-D23
Termination
D23
D0
D7
D14
D21
MM.1S
subcut.
injection
NSG
mice
E
D23 Tumour weights
Tumour weight (gr)
Control
T417
F
Control
T417
Body weight (gr)
(gain/loss relat. to Day0)
Time (days)
G
Cells
SSC-A
FSC-A
Singlets
FSC-W
Human
(HLA+)
HLA-ABC
Live
Propidium Iodide
Annexin V
Cell cycle
Control
T417
G0/1
S
G2/M
Count
Hoechst
H
in vivo
Live cells (%)
(HLA+/AnnexinV-PI-/Hoechst+)
G2/M
S
G0/1
Control
T417
I
X1
Live cells (%)
(AnnexinV-PI-/Hoechst+)
J
PBBC
K
PBX1
FOXM1
NEK2
E2F2
AMO-WT
AMO-BZ
Undet.
Rel. expression
(T417 vs control)
L
Bortezomib (nM)
AMO-WT
AMO-BZ
T417 (μM)
Combined effect
antagonistic
additive
synergistic
M
X1
X3

**Supplementary Fig. S7 related to Fig. 6. A novel PBX1 small-molecule inhibitor for selective targeting of chr1q-amp cancer.**

**(A)** Statistical overview (Hazard ratios: HR, P-values: P) of overall survival analysis between patient arms with high versus low PBX1 activation across 8 solid tumour patient cohorts.

**(B)** Number of chr1q copies across multiple myeloma (blue), breast (red), ovarian (green), lung (purple) and (cyan) cancer cell lines. Genetic amplification of chr1q was determined by FISH (for MM1.S, NCU.MM1, OPM2, U266, MCF-7, LTED and A2780) or by using the *PBX1* CNV scores from the Depmap database (AMO.1, OVCAR-3, A549, H69AR and SNB-75)

**(C)** Sensitivity of multiple myeloma (MM1.S, NCU.MM1) and ovarian cancer (OVCAR-3, A2780) cell lines to the active PBX1 inhibitor T417 (PBXi) or the inactive analogue DHP52 (PBXi_mock). Non-linear fitting was performed as previously described. One-way ANOVA with post-hoc multiple comparisons test was used to compare PBXi versus PBXi_mock-treated cells for each cell line. Error bars show standard errors of mean. ***: $P<0.001$; ****: $P<0.0001$.

**(D)** Schematic diagram of *in vivo* experiment plan using a MM xenograft model. In brief, approximately $10x10^6$ MM1.S cells were injected subcutaneously in 6 male and 6 female NSG mice (D0). When all tumours reached a measurable size (D7), mice were randomized and split into two arms: control (vehicle) and T417 (10 mg/kg/injection) arm, 3 male and 3 female mice per arm. Mice were treated with vehicle or T417 via intraperitoneal (I.P.) route with a 4 days on – 3 days off regime for a total of 10 courses (D8-D11, D15-D18, D22-D23). Body weights and tumour sizes were measured daily. Experiment was terminated when tumours reached the maximum allowed size (D23); all tumours were extracted and analysed on the same day.

**(E)** Tumour weights of control and T417-treated mice upon termination (D23). Comparison was performed using a Mann-Whitney test. Error bars display the standard errors of mean. ****: $P<0.0001$.

**(F)** Timecourse measurements of mouse body weights for control (black) and T417 (orange) treatment arms. Values are shown as normalized to Day0.

**(G)** Gating strategy for cell cycle analysis of extracted tumours on termination day (D23). FACS plots illustrate an example of tumour Cells/Singlets/HLA+/Live population profiles in control- and T417-treated mice.

**(H-J)** Cell cycle analysis of **(H)** isolated tumours (control vs T417 arms) on termination date (*in vivo*), **(I)** primary chr1-amp MM cells (X1), and **(J)** *in-vitro* cultured normal donor primary peripheral blood B cells (PBBCs) 48h after treatment with control (1% DMSO) or T417 (20µM). Three independent experiments were performed for PBBCs and one for X1. Statistical analysis was performed using a two-way ANOVA with post-hoc multiple comparisons test.

**(K)** Relative expression of *PBX1, FOXM1, NEK2* and *E2F2* in AMO.1-WT and AMO.1-BZ cells 20h after treatment with 20µM T417, relative to 1% DMSO (control) treatment.

**(L , M)** Analysis of T417-Bortezomib combined effect on viability of parental (AMO.1-WT), bortezomib-resistant MM cell lines (AMO.1-BZ), as well as of primary chr1q-amp MM cells (X1, X3). Heatmap illustrates the combination index (CI) scores simulated by CompuSyn software. (-), antagonistic; 1, additive; (+), synergistic effect.
